# Transcriptogram analysis reveals relationship between viral titer and gene sets responses during Corona-virus infection

**DOI:** 10.1101/2020.06.16.155267

**Authors:** Rita M.C. de Almeida, Gilberto L. Thomas, James A. Glazier

**Author notes:** **Corresponding author: Prof. Rita M.C. de Almeida, Instituto de Física, Universidade Federal do Rio Grande do Sul, Porto Alegre, RS, Brazil, / 55 51 982090106**.

## Abstract

To understand the difference between benign and severe outcomes after Coronavirus infection, we urgently need ways to clarify and quantify the time course of tissue and immune responses. Here we re-analyze 72-hour time-series microarrays generated in 2013 by Sims and collaborators for SARS-CoV-1 *in vitro* infection of a human lung epithelial cell line. Transcriptograms, a Bioinformatics tool to analyze genome-wide gene expression data, allow us to define an appropriate context-dependent threshold for mechanistic relevance of gene differential expression. Without knowing in advance which genes are relevant, classical analyses detect <u>every</u> gene with statistically-significant differential expression, leaving us with too many genes and hypotheses to be useful. Using a Transcriptogram-based top-down approach, we identified three major, differentially-expressed gene sets comprising 219 mainly immune-response-related genes. We identified timescales for alterations in mitochondrial activity, signaling and transcription regulation of the innate and adaptive immune systems and their relationship to viral titer. At the individual-gene level, EGR3 was significantly upregulated in infected cells. Similar activation in T-cells and fibroblasts in infected lung could explain the T-cell anergy and eventual fibrosis seen in SARS-CoV-1 infection. The methods can be applied to RNA data sets for SARS-CoV-2 to investigate the origin of differential responses in different tissue types, or due to immune or preexisting conditions or to compare cell culture, organoid culture, animal models, and human-derived samples.

## Introduction

Severe respiratory syndromes during the two previous major outbreaks of lethal Coronavirus, SARS-CoV-1 in 2003 [1] and Middle-Eastern Respiratory Syndrome (*MERS*) in 2012 (for a Review, see [2] and references therein), as well as the current SARS-CoV-2 pandemic, often result from dysfunctional immune responses triggered by the interaction of the host immune system with the virus [3,4]. While strong immune responses are essential to contain and clear viral infection, excessive inflammation may damage tissues, delay tissue healing after viral clearance, and lead to acute inflammatory responses and/or sepsis. In the case of SARS-CoV-2, the degree and severity of immune-response pathologies differ greatly between individuals (https://globalhealth5050.org/covid19/). Because of the complexity of the many patterns of response to SARS-CoV-2, we critically need ways to identify important biological mechanisms which act at different phases of infection and allow us to reliably identify differences in pathway and gene activity between individual patients, tissues within patients, individuals with pre- exiting conditions, sex and ethnic differences and age. The immune system is complex, sensitive and dynamic, with a delicate balance of triggers, high-gain feed-back loops, and complex interactions between its many agents, complicating interpretation of experimental measurements of immune-response components and the origins of their variation between individuals. In this case, for diagnostic, prognostic, and therapeutic purposes, detailed mathematical models of patient-specific immune responses might help us understand the range of possible immune responses, and how they depend on patient-specific variables, ranging from initial exposure level and coinfections, to age, sex, preexisting conditions and medications, *etc*. Furthermore, in serious cases, COVID-19 symptoms may also include blood and vascular disruption, meaning that the co-activation of other pathways with deleterious effects may play an important role in disease outcomes [5].

Both constructing mathematical models of a complex system like the human immune response and validating such models sufficiently for use to propose therapies or assist with diagnoses or prognoses, requires integration of extensive data from *in vitro*, organoid and animal experiments with the more limited clinical observations in humans. Acute inflammatory responses lead to dramatic and rapid changes in expression of large numbers of genes, requiring extensive transcriptome analyses to interpret. For construction and validation of immune-response models, qualitative information is insufficient; we also need specific quantitative information on the time course of immune response and its relationship to viral titer.

Statistical analyses of transcriptome data are generally classified as either *bottom-up*, starting by identifying differentially-expressed genes, clustering them into differentially-expressed pathways and then describing the biological functions these pathways alter, or *top-down*, starting by identifying altered biological functions, then refining the analysis to hierarchically discover the relevant differently-expressed pathways and then genes. In cases of immune-system response to viral infection, where changes in gene expression are genome-wide, top-down approaches may be more practical, since the large number of differentially-expressed genes can be overwhelming to analyze and understand using bottom-up techniques.

RNA-Seq or microarray transcriptomes are affected by many sources of variability, including differences in experimental techniques, biological differences between apparently similar samples, and other confounding variables within samples, like the effect of cell-cycle phase. Our Transcriptogram method to quantify whole-genome-level expression changes reduces noise and enhances signal-to-noise ratio in transcriptome analyses, increasing the power of statistical tests to identify significantly affected pathways and timescales [6]. Transcriptograms provide a high-level visualization of significant changes in gene expression and have proved useful in identifying relationships between pathways in fungi [7,8], plants 9,10, and humans [11,12,6]. The Transcriptogramer software tool is freely available for download at https://lief.if.ufrgs.br/pub/biosoftwares/transcriptogramer/ and has a Bioconductor application [13].

Here, as a pattern for future Transcriptogram analyses of SARS-CoV-2 data and to illustrate the power of the method in quantifying the detailed and complex temporal pattern of immune response to viral infection in cell culture, we present Transcriptogram analyses for SARS-CoV-1 time-series data sets of Sims *et al*. [14]. Sims *et al*. [14] infected cultures of a clonal population of Calu3 2B4 cells, a lung adenocarcinoma cell line isolated from the pleural effusion of a 25-year-old Caucasian male, sorted for high expression of the enzyme ACE-2, a major cellular receptor for SARS-CoV-1 (and SARS-CoV-2). They inoculated cultures with either a wild type SARS-CoV-1 virus (*WT* samples) or a mutant SARS-CoV-1 strain (*DORF6* samples) that does not express the accessory protein ORF6 at high concentration (a multiplicity of infection *MOI* of 5), so that the probability of cell contamination in the culture approached 1. As controls, they also inoculated cultures with a sterile solution (*Mock* samples). After inoculation, they incubated the cultures at 37°C for 40 min, then changed their medium. They then harvested samples for microarray assays in triplicate at times they labeled 0 *h*, 3 *h*, 7 *h*, 12 *h*, 24 *h*, 30 *h*, 36 *h*, 48 *h*, 54 *h*, 60 *h*, and 72 *h*. Because they did not report the time for the medium change or the time between inoculation and initial harvest, their data lack a consistent time-0 data set and all time labels refer to the time after the first RNA harvest. As a result, even at 0 *h*, expression in the infected and control cultures differs (see below). We analyzed these data because of the short time intervals between samples at early times, which are critical to understanding the rapid changes occurring in tissue response to viral infection, and because of the relatively long duration of the experiment. The experiments also have matched-time controls in triplicate at all time points. Sims *et al*. [14] made their data available at Gene Expression Omnibus (*GEO*) under accession number GSE33267 (https://www.ncbi.nlm.nih.gov/geo) and we used these data for our analyses. More details on the experiments are available in Supplementary Information online, section 1.

Sims *et al*. focused their analyses on the role of ORF6 in the immune response, examining the differences between the *WT* and *DORF6* time series [14]. Here, we focus on large-scale and single-gene transcriptomic changes caused by the *WT* virus *w*.*r*.*t*. the control. Our analyses confirm that gene expression changes massively within 24 *h*, but we also identified relevant responses before 7 *h* and complex temporal changes in expression throughout the time-course of the experiment. Our analyses identify specific additional significant changes in expression in different pathways and individual immune-related genes at 12 *h*, 36 *h* and 54 *h*. We identified 219 genes with differential expression at some point of the time sequence for *WT* samples *w*.*r*.*t* the control, classified these genes in three clusters by their covariance, and monitored the evolution of the mean differential expression for each cluster. The results suggested hypotheses regarding the cellular response to the virus in these experiments, as we discuss below. We also examined these clusters’ mean differential expression for the mutant virus strain samples and present these results in the Supplemetary Information online. To illustrate the potential of our method, we selected 4 genes with large expression differences *w*.*r*.*t*. to controls for further scrutiny, EGR3, TWIST1, JUN, and TNFAIP3, all related to immune response. To validate our findings, we also examined a pair of genes, HSD11B1 and HSD11B2, with known associated effector action on Cortisol/Cortisone balance.

We also remark that the cellular immune response accompanies genome-wide changes in gene expression, yielding too many genes with statistically-significant differential expression to be helpful in identifying critical pathways. Separating the very many genes with statistically-significant changes in expression from the many fewer genes which are biologically relevant to the immune response can be compared to finding a needle in a haystack. Transcriptogram analysis acts as a filter based on gene function, reducing the number of genes of interest to a tractable set and suggesting shared mechanistic functions for observed gene expression patterns.

## Statistical Methods

### Overview and analyses pipeline

We performed a Transcriptogram-based [8,15] top-down analysis of whole-genome transcriptome time-series for human epithelial cells cultures, comparing cultures inoculated with either a control (*Mock*), a wild type SARS-CoV-1-containing (*WT*) solution, or a mutant SARS-CoV-1-containing (*DORF6*) solution (GEO accession number GSE33267 (http://www.ncbi.nlm.nih.gov/geo)). We first assessed patterns of expression change shared by large numbers of genes, then considered smaller gene sets which had strongly covariant temporal signatures, and finally examined single genes whose variance was statistically significant withing these sets. At each stage, we filtered the data based on the statistical significance of the subseries variability. The next section briefly discusses the Transcriptogram method and relevant parameters. We focused our analyses on the cellular response to *WT* virus inoculation. We present the data and Transcriptograms for the *Mock* and Wild Type (*WT*) virus-strain time courses in the main text and the Transcriptograms for DORF6 in the Supplementary Information online. We indicate the gene sets we focus on, and provide a genome-wide visualization of the main patterns of the time evolution for the covariance-clustered gene data, comparing the *WT*-infected and non-infected samples. We identified 219 genes whose time courses for the *WT*-samples show fold-changes larger than two compared to their pair-matched *Mock* samples at at least one time point. We clustered these genes by time-course similarity and identified 3 clusters. We determined the time evolution for each cluster’s mean expression for both *WT* and DORF6 samples. These clusters show complex non-monotonic time courses, which we compare to the viral titer. From the changes in behaviors presented by the gene clusters’ mean expression, we infer the typical patterns and timescales for Calu3 2B4 epithelial-cell immune response to SARS-CoV-1 infection, correlate these timescales with viral titer and identify single genes that may be possible targets for therapy development.

### Transcriptograms

*Transcriptograms* are expression profiles, obtained by running a window average for expression levels of multiple genes, previously organized in an ordered gene list, representing the whole human genome. Here we consider windows of radius 30, that is, intervals around a given gene position including 30 genes to its left and 30 genes to its right in the ordered gene list. Averaging over these intervals for each gene in the list produces a smoothed mean expression profile. We generate the ordered gene list by first filtering gene products that share at least one association as inferred from the STRING Protein-Protein Interaction database with confidence scores of 800 or better [11]. The gene-list ordering clusters genes by their biological function as defined in the STRING database. STRING uses seven different methods to infer whether two proteins are associated, ranging from physical interactions, information retrieved from (reliable) data bases like HUGO, to co-expression, experiments, *etc*.. STRING also provides a confidence score for the inference, given by the fraction of inferences of each method that predicts that the two proteins participating in the same KEGG metabolic pathway. We used an overall STRING score of 0.800 as our threshold for accepting an interaction. The STRING information we use is more far-reaching than physical interactions, and does not rely on single experiments, but on knowledge about protein-protein association that integrates contributions from an extensive scientific community.

The ordered gene list we use for this analysis is available as an additional file AF1 in Supplementary Information online. Ref. [11] explains the construction of the gene list in detail. We then apply this ordered list of genes to analyze gene expression data from micro-arrays or RNA-Seq experiments. Because the list clusters genes by attributed function, the running window averages expression levels over genes believed to participate in the same or similar biological functions.

One major problem in detecting differential gene expression in microarray or RNA-Seq experiments is with the high variance of the data, that can result from measurement noise or confounding variables that were not explicitly controlled for. Ref. [6] shows that Transcriptograms can reduce the variance of gene expression measurements and enhance the power of statistical tests when comparing gene expression levels between gene samples. We characterize the ordered genes by projecting onto the gene list selected biological Gene Ontology (*GO*) terms or KEGG pathways, to associate regions of the list with key biological mechanisms.

Fig. 1 shows term-enrichment profiles projected on the ordered list, obtained for selected KEGG pathways and Gene Ontology: Biological function terms (*GO:BP*). The gene list we use comprises 9684 genes, representing those genes whose products participate in at least one Protein-Protein Interaction (with a score of 800 or better) as listed in STRING. The horizontal axis (intrinsically numbered by gene position from 1 to 9684) has been rescaled to fit the interval [0,1]. At each position in the gene list, represented by the horizontal axis, we plot the fraction of genes within a window of radius 30 genes around that position associated with a specific term or pathway. A profile value near 1 means that almost all 61 genes in that interval link to the term. Moving from left to right, we observe successive enrichment of terms associated with specific biological functions: at the far left, we see enrichment linked to RNA processing and metabolism, then enrichment related to the cell cycle, followed by cell differentiation, the actin cytoskeleton and immune systems. Further to the right, we see enrichment for signaling pathways associated with secretion, ECM receptors and finally, energy metabolism. Consequently, a running window average of expression data over this ordering, averages the expression of genes linked to the same or similar biological functions.

**Fig 1.**
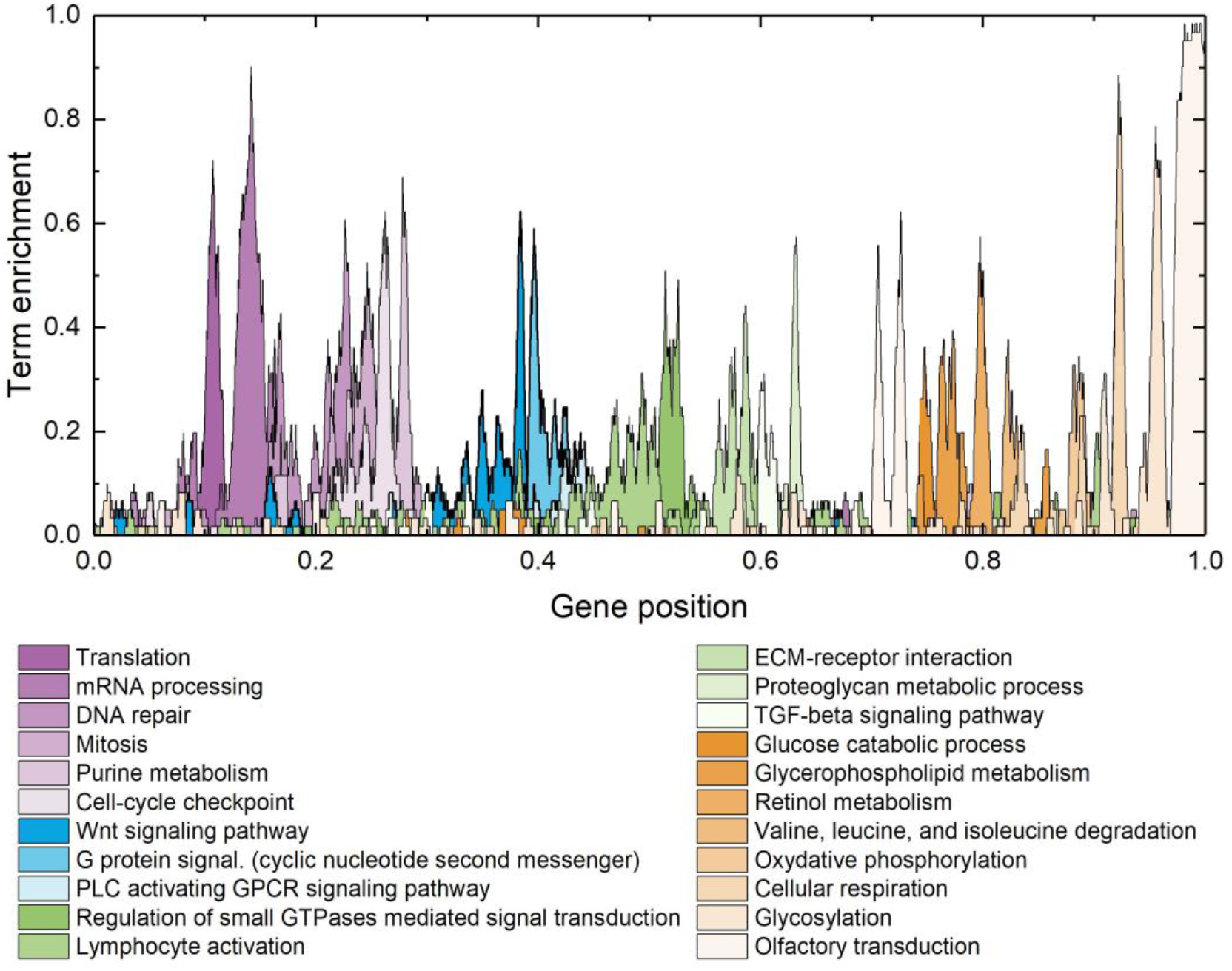
Gene list and enrichment of terms related to critical biological functions as a function of position in the list. From left to right, in shades of purple, the list is enriched with genes associated with translation and mRNA processing then pathways linked to the cell cycle. Next, in shades of blue, genes associated with cell differentiation and, in shades of green, genes associated with immune response, cytokine production and interaction with the extra-cellular matrix (ECM). Finally, shades of orange denote genes associated with energy metabolism.

### Normalization check using Transcriptograms

Transcriptograms provide a powerful test for sample normalization by revealing undesired variable offsets in expression levels between samples. We ensure sample normalization as follows: we plotted the Transcriptograms of radius 30 for the normalized data available in the GEO database (shown in Fig. S 1 in the Supplementary Information online) and verified that each sample set shows offsets in its mean in relation to the other sample sets. We then re-normalized the expressions levels for each sample data set to set its mean expression to 1. Fig. S 1 shows the resulting renormalized, single-sample Transcriptograms.

### Relative Transcriptograms

We obtain *relative Transcriptograms* by dividing the Transcriptogram profile values at each point in the ordered gene list by the Transcriptogram profile value for a control sample at the same position in the list.

### Differential Transcriptograms

We obtain *differential Transcriptograms* between **two time-series** by obtaining the relative Transcriptogram for the *WT* samples at a given time point *w*.*r*.*t*. time-matched *Mock* samples. In a time-series for differential Transcriptograms, the control sample differs for each time point.

### Term Enrichment

We determined term enrichment for the gene sets consisting of the genes in a given interval of the ordered gene list using the Term Enrichment Panther Service, on the Amigo 2 home page (https://amigo.geneontology.org/amigo) [16,17,18].

### Covariance matrix

For each gene *i* from a gene set with elements (*i* = 1, …, *N*) we define the *differential expression e*_*i*_(*t*) as:

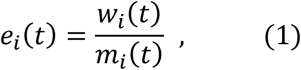

where *t* represents a time in the time series and *w*_*i*_(*t*) and *m*_*i*_(*t*) are the averages over replicates for the gene expression values from, respectively, the *WT* or Mock transcriptomes’ normalized datasets.

We define the *covariance matrix C*_*ij*_ as:

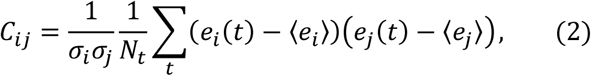

where *N*_*t*_ is the number of the experiment time points (here *N*_*t*_ = 11) and ⟨ *e*_*i*_ ⟩ is the time average of the differential expression of the *i*-th gene, that is:

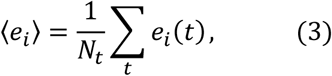

and σ_*i*_ is the standard deviation calculated as:

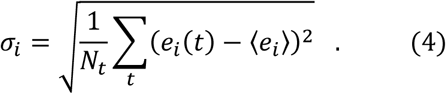

## Results

We start our analysis of expression profiles for *WT* and *Mock* samples at different times by generating relative Transcriptograms as described in the Statistical Methods section, taking as the control the Transcriptogram for the *Mock* sample at time 0 *h* (which is the time of the first RNA harvest, at least 40 minutes after inoculation). Fig. 2 shows the relative Transcriptograms at different times for *Mock* (blue lines) and *WT* (red lines) samples. The relative Transcriptogram for the control expression levels (*Mock* samples at 0 *h*) appears as a black horizontal line. We also plot the relative Transcriptogram standard errors (due to the variance among replicates) for each point of the ordering: these errors are represented by gray, light red, and cyan shading around, respectively, the black, red and blue lines. The Transcriptogram’s window average reduces the variance between replicates, so the error bars are barely visible.

**Fig 2.**
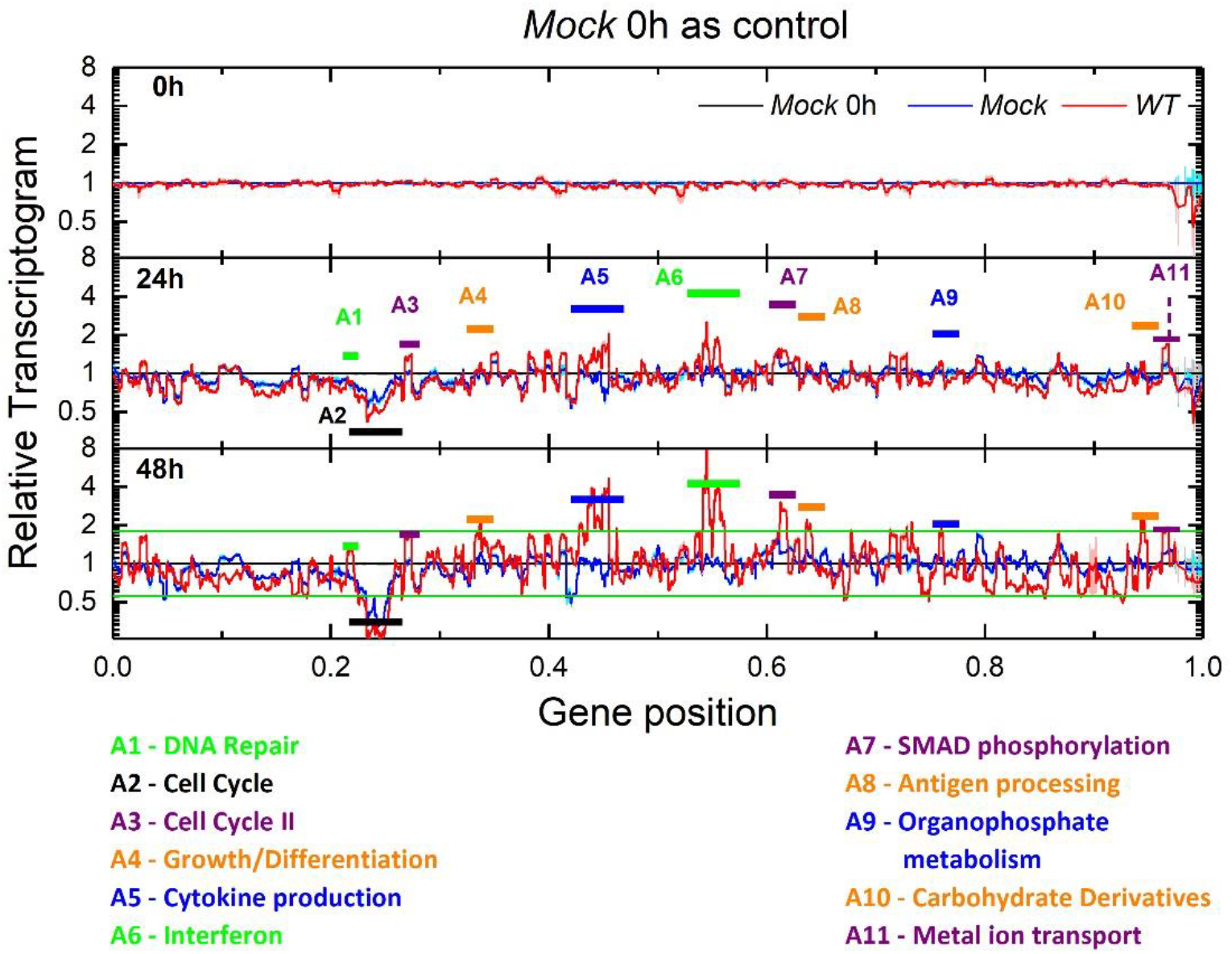
Relative Transcriptograms of radius 30 for Mock and WT samples, using the Mock sample’s expression at t = 0 has the control. The labeled time is the experimental time after the first RNA harvest (over 40 minutes after inoculation). Vertical axes are on a log_2_ scale. Green horizontal lines in the bottom panel correspond to 9/5 and 5/9 fold changes. Black horizontal lines represent the control sample (Mock) expression. Red and blue lines represent the relative Transcriptograms for, respectively, WT and Mock samples. Gray, light red and cyan shading indicate the standard errors of the respective relative Transcriptograms. We identify 11 intervals, indicated by the horizontal color bars, where the red line differs from the control by more than 9/5 at 48 hor after, except for peak A1. (See the panel for the complete time series in Supplementary Information online, Fig. S 6).

Fig. 2 presents the relative Transcriptogram profiles at 0 *h*, 24 *h* and 48 *h* after first RNA harvest. The top panel shows that at *t* = 0 *h* (approximately 40 minutes after the viral inoculation and washout at the start of the experiment) the *Mock* and *WT* samples are similar, although in some regions the errors do not overlap (around gene positions 0.41 or 0.53, for example), indicating that cells are already responding to viral infection at the time of first RNA sampling. Fig. 2 shows that at 24 *h* and 48 *h*, the relative Transcriptogram profiles of both *Mock* and WT samples differ significantly from the *Mock*-sample mean profile at 0 *h*. The direction of deviations at a given gene location at 48 *h* is typically the same as that at 24 *h*, but of greater amplitude. To identify intervals in the relative Transcriptograms with significant expression variations, we define and consecutively label contiguous regions along the horizontal axis with values larger than 9/5-fold changes in the *WT* relative Transcriptogram values at 48 *h* (labels A1, A2, …,A11). From the many terms enriching each region (see Methods section) we select a representative term as a label, based on the number of genes associated to that term in that region.

Fig. S 6 in the Supplementary Information online shows equivalent panels for all time points (0 *h*, 3 *h*, 7 *h*, 12 *h*, 24 *h*, 30 *h*, 36 *h*, 48 *h*, 54 *h*, 60 *h*, 72 *h*). While our method does not seek to identify genes related to immune response, most of the bands having statistically significant different alterations correspond to regions enriched with genes participating in pathways linked to the immune response. Genes linked to the cell-cycle I region (interval A2) are depressed in both *Mock* samples and *WT* samples, probably reflecting contact inhibition of proliferation in confluent *in vitro* cultures. However, Fig. S 6 in the Supplementary Information online also shows that at 54 *hWT* samples show some expression recovery of genes linked to the cell cycle. This recovery may reflect the onset of cell cycle after the death of some infected cells, which reduces contact inhibition of proliferation, or another tissue-recover mechanism. Changes in expression across multiple functional bands of the relative Transcriptogram appear at 24 *h*. These bands stay fixed in width but increase in amplitude until 54 *h*, after which they slowly decrease in amplitude. For more details on the changes in each band, refer to Fig. S 6 and movies SM1 and SM2 in the Supplementary Information online.

The important messages in Fig. 2 and Fig. S 6 and the animation of the time changes of the relative Transcriptograms in movies SM1 and SM2 are that: i) major changes in band expression start after 12 *h*; ii) the bands of expression change in amplitude but not in width, reflecting their correspondence with changes in activity of specific biological mechanisms; and iii) gene expression in the control samples also changes in time, because cell state changes in culture conditions, even in the absence of infection.

To distinguish cell-culture effects, which affect both *WT* and control cultures, from the effects of infection, we considered time-matched *Mock* expression profiles as controls for the *WT* expression profiles. We define the *differential* Transcriptogram profile as the ratio of the *WT* transcriptogram value at a given time and gene position to the time-matched *Mock* transcriptogram value at the same gene position. Using a time-matched control helps reduce the signal from tissue-culture effects common to both *WT* and control samples and accentuates specific differential infection effects. Differential profiles do not show changes in expression of cell-cycle-related genes, for example, since both control and *WT* expression change in the same way in time. Fig. 3 shows differential profiles as violet lines, with the light violet shading showing the standard error. The horizontal black line shows the control differential expression profile for the *Mock* sample at the corresponding time. Fig. 3 presents the differential profiles (*WT* (t)/*Mock* (t)) at three time points, Fig. S 7 in the Supplementary Information online presents the differential profiles for all time points and Supplementary Movie SM3 animates these time changes.

**Fig 3.**
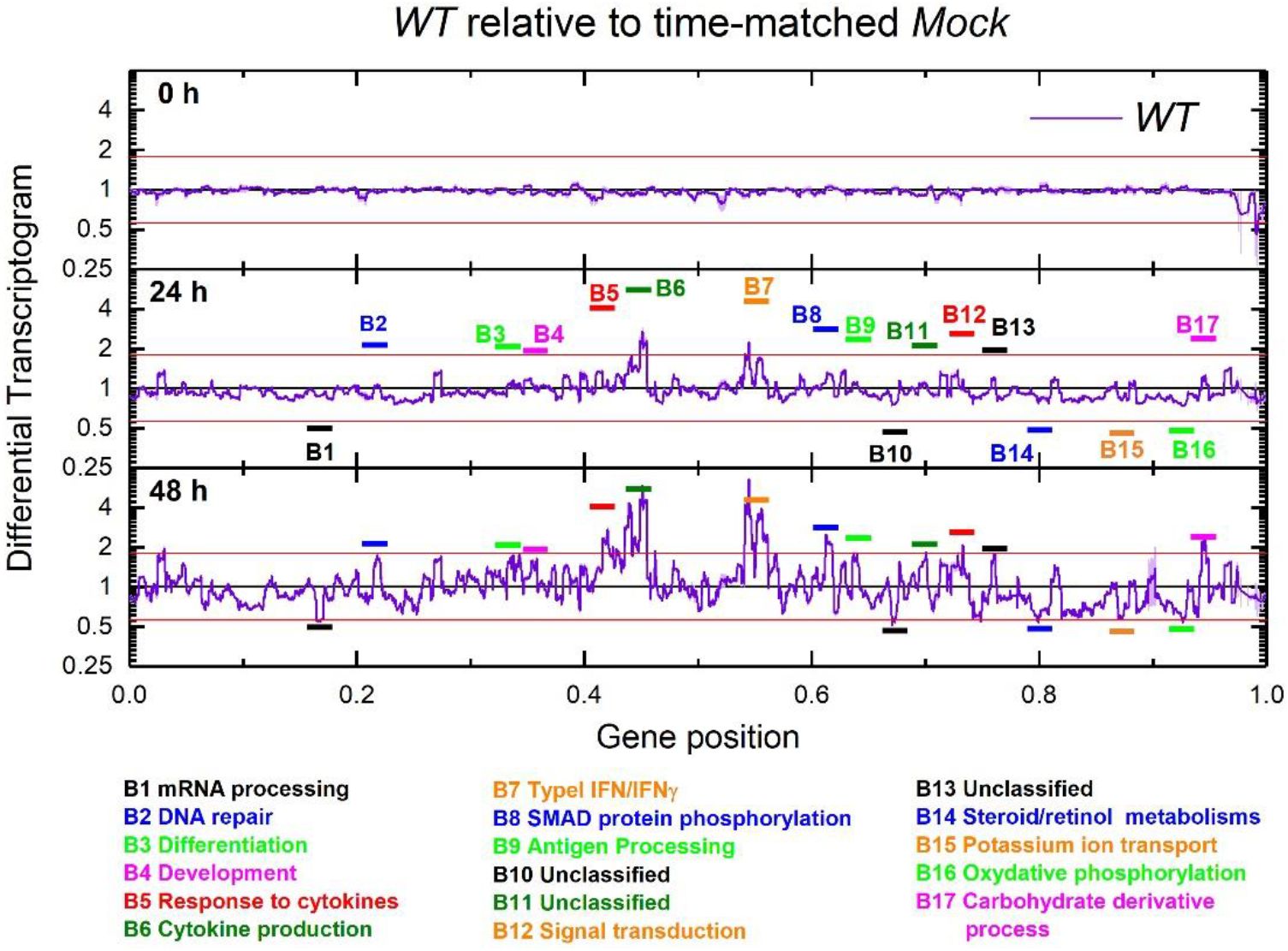
Differential Transcriptograms WT(t)/Mock(t) (radius 30). Time is time after the first RNA harvest. Vertical axes are on a log_2_ scale. Black horizontal lines represent the control sample Mock(t)/Mock(t). Violet lines are the differential Transcriptograms for WT(t)/Mock(t). Light violet shading indicates standard errors for WT Transcriptograms. We identify 17 bands where the violet line differs from the control more than 9/5-fold at 24 h. The horizontal red lines denote the 9/5-fold and 5/9-fold lines in all panels.

Differential Transcriptogram profiles show noticeable alterations after 12 *h*. As before, we associate the most altered bands to biological functions using DAVID tools [19] at https://david.ncifcrf.gov/tools.jsp and label them accordingly (see below). As in Fig. 2, the altered bands remain constant in width, but change in amplitude.

We next consider the time evolution of the differential expression (*WT* (*t*)/*Mock* (*t*)) of the 590 individual genes that participate in the 17 bands identified as significantly differentially expressed in the differential Transcriptograms in Fig. 3 (where we define a significant change as either larger than 9/5 or less than 5/9). Additional file AF2 in the Supplementary Materials online presents these results as an Excel file containing the gene names and plots for each gene’s relative and differential expression evolution, with brief information on each gene. Among these 590 genes, we found 219 for which the expression *w*_*i*_(*t*) at one or more time points *t* differed from the time-matched control *m*_*i*_(*t*) more than two fold (*i*.*e. w*_*i*_(*t*) > 2*m*_*i*_(*t*) or *w*_*i*_(*t*) < 0 5 *m*_*i*_(*t*)). Our significance limit is higher because here we are considering single-gene differential expression, rather than the differential Transcriptogram values, which are averages over the expression of neighboring genes in the list.

The Transcriptogram analyses identify 219 genes in differentially-expressed Transcriptogram bands which are also individually differentially expressed relative to their time-matched *Mock* samples. This gene set comprises the genes that respond more intensely to virus inoculation. To identify their associated biological functions, we used the Over Representation test by Panther, available on the Gene Ontology-Amigo home page (https://amigo.geneontology.org/amigo) to find the Reactome Pathways that enrich this set of 219 genes. Among others, we find that 51 of these genes participate in “Cytokine signaling in immune system,” 36 participate in “Innate immune system,” 18 participate in “Toll-like-receptor cascade,” 15 participate in “Interleukin 4 and Interleukin 13 signaling” and 8 genes are participate in “TNFR2 non-canonical NF-κB pathways.” Additional file AF3 in the Supplementary Materials online gives the complete list of over-represented Reactome Pathways for the 219 genes (P<0.05, Buonferroni corrected). Of the 219 genes, 35 have not been classified as forming a representative set for any Reactome Pathway. Thus 84% of the 219 significantly variant genes participate in pathways either directly or indirectly involved in components of the immune response.

Proceeding with our top-down strategy, we look for temporal patterns in the differential expression of single genes (not Transcriptogram values). To find genes with similar patterns of temporal evolution, we first calculated the covariance matrix for the differential expression time series of the 219 genes (see Statistical methods section). When two genes have the same pattern of time change of differential expression, their temporal covariance approaches one. Using the covariance matrix, we ordered genes into covariance clusters (Fig. 4 A). We identify 3 clusters, A, B and C. As discussed previously, the ordering of the gene list is based on biological function attributed to the genes. Now, the covariance matrix clusters genes by the similarity of their differential expression time series. The activation a pathway associated with one biological function may lead to activation of genes associated with different biological functions, leading to temporal correlations in their differential expression patterns. Also, different genes within a pathway may activate with different time patterns, reducing their temporal covariance. Consequently, we do not expect the covariance matrix clusters to directly correspond to the differential transcriptogram bands in Fig.3. Instead, Clusters A to C are covariant gene sets for the genes which both Transcriptogram and single-gene analysis identified as significantly differentially expressed in the present experiment. Additional file AF2 in the Supplementary Materials online is an Excel file containing a separate worksheet for each of the 17 bands which the Transcriptogram analysis identified as significantly differentially expressed, as shown in Fig. 3. For each of these bands, file AF2 presents plots of the relative and differential expression of the 219 genes, in different colors, depending on the time-series pattern they follow. All bands have genes which belong to all covariant clusters, showing that proximity in the ordered list does not correlate with covariant cluster identity.

**Fig 4.**
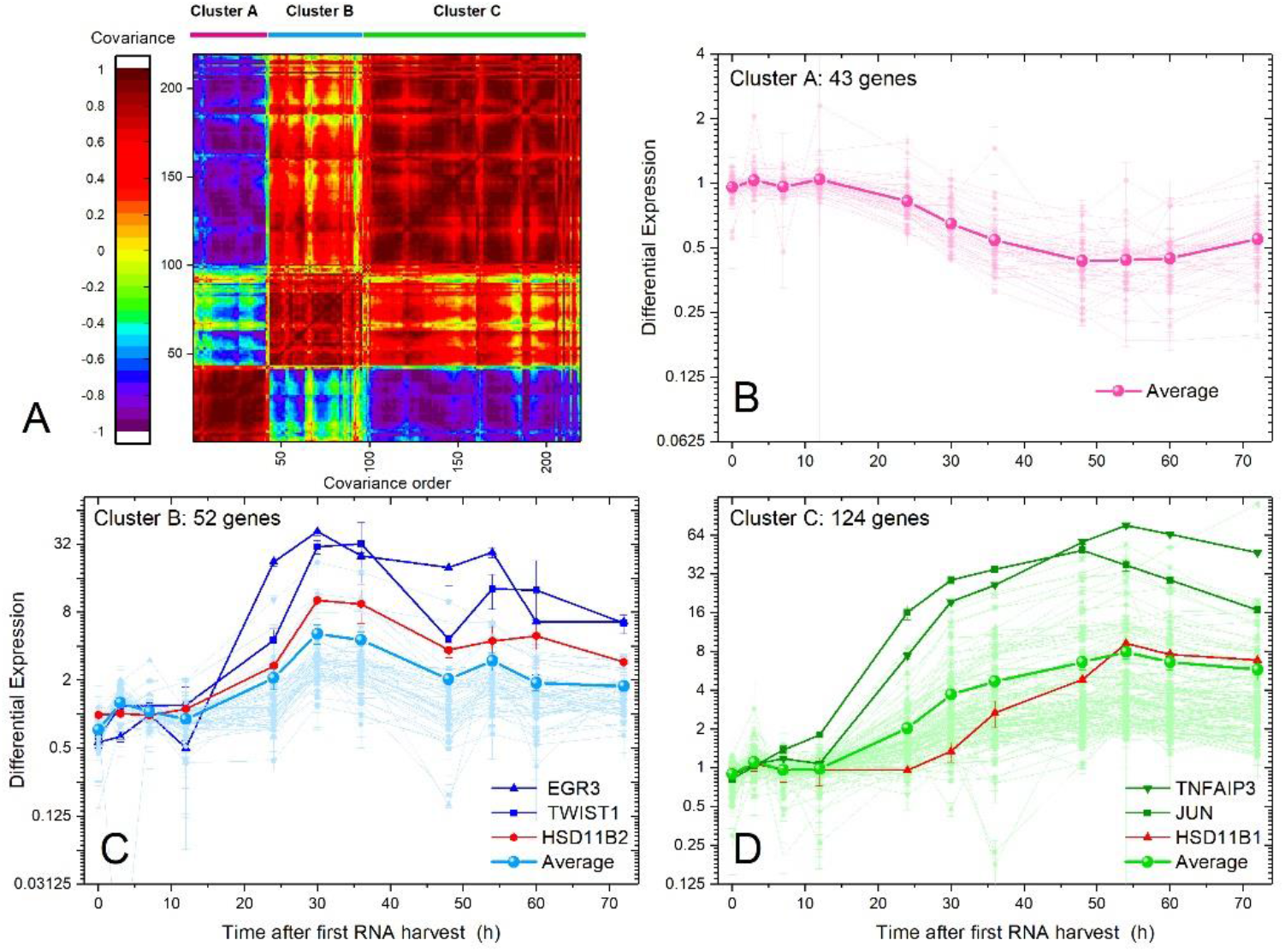
Covariance matrix for the time changes of differential expression (time-matched control) for 219 genes selected from the significantly differentially expressed Transcriptogram bands identified in Fig. 3. We find 3 major covariant clusters. Table 1 lists the genes in each cluster. Figs. 4 B, C, and D show the time-series for the differential expression for each gene in each cluster and the averaged differential expression time-series for each cluster. Selected genes in each cluster are highlighted. The control for each gene at each time point is the Mock sample expression of the same gene at a matched time. Expression is for individual genes, not Transcriptogram averages over the neighbors in the ordered gene list. Clusters A (43 genes), B (52 genes), and C (124 genes) have highly covariant differential expression time series. Additional file AF2, in the Supplementary Materials online shows the individual relative expression time series for each gene in full detail.

### Error! Reference source not found

(B to C) also present the time series for the differential expression of each gene in each cluster, together with the cluster average of these values. Within clusters A, B, and C the genes have similar patterns of change of differential expression in time. Fig. 5 summarizes the averaged temporal patterns of differential gene expression of the clusters.

**Table 1.**
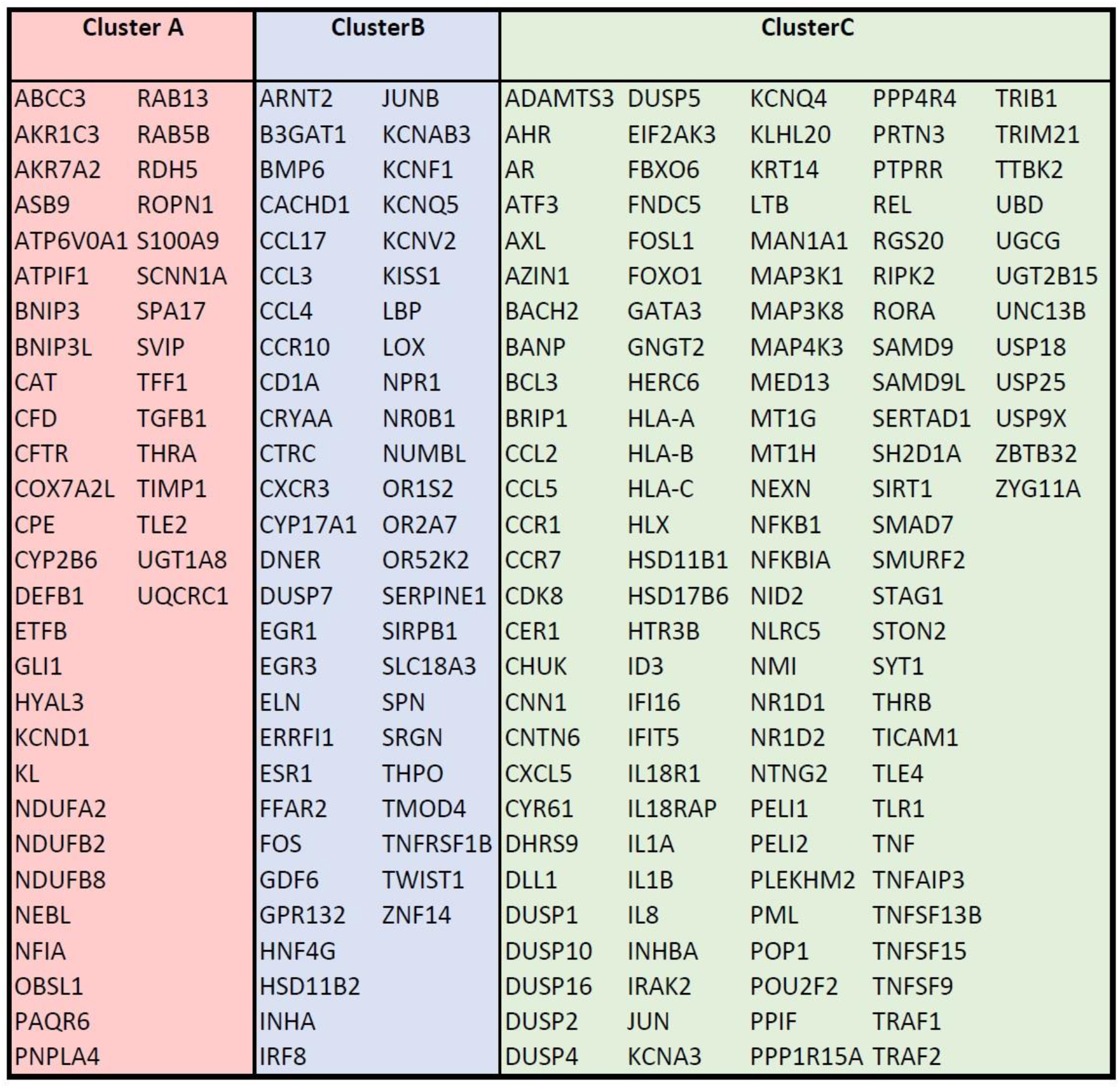
List of genes in each covariant cluster (Fig. 4). Cluster A mostly relates to energy metabolism, Cluster B is enriched with genes related to cytokine production, and Cluster C involves genes related to cell response to stimuli, including cytokines.

**Table 2.** P values comparing the mean expression of Clusters A, B, and C for the whole time series. Columns on the left compare WT to Control samples, and columns on the right compare clusters, two by two, for the time-matched differential expression represented in Fig. 5.

| Time after<br>first RNA<br>harvest (h) | P-values |  |  |  |  |  |
| --- | --- | --- | --- | --- | --- | --- |
|  | WT and Mock matched time samples |  |  | Between Clusters |  |  |
|  | Cluster A | Cluster B | Cluster C | Clusters A<br>and B | Cluster A<br>and C | Cluster B<br>and C |
| 0 | 5.50E-02 | 2.19E-04 | 8.59E-06 | 1.35E-09 | 6.16E-03 | 1.85E-06 |
| 3 | 1.78E-03 | 4.78E-01 | 6.29E-01 | 1.97E-04 | 7.21E-02 | 6.59E-03 |
| 7 | 2.16E-01 | 3.35E-04 | 8.26E-01 | 9.99E-02 | 9.21E-01 | 1.05E-01 |
| 12 | 9.64E-01 | 3.53E-02 | 2.89E-01 | 1.18E-02 | 1.22E-01 | 8.38E-02 |
| 24 | 9.35E-03 | 3.71E-03 | 4.89E-04 | 6.90E-03 | 6.77E-11 | 8.96E-01 |
| 30 | 1.29E-06 | 2.29E-02 | 1.31E-03 | 4.25E-05 | 4.13E-13 | 1.79E-01 |
| 36 | 1.15E-05 | 4.60E-03 | 6.17E-04 | 9.31E-06 | 9.39E-14 | 8.70E-01 |
| 48 | 6.17E-04 | 3.11E-04 | 1.15E-06 | 3.08E-04 | 1.52E-11 | 2.13E-06 |
| 54 | 7.79E-04 | 6.93E-05 | 1.83E-04 | 2.51E-05 | 6.80E-12 | 1.83E-05 |
| 60 | 3.14E-03 | 8.51E-03 | 7.10E-04 | 1.09E-06 | 1.59E-11 | 2.22E-07 |
| 72 | 1.62E-04 | 1.34E-03 | 1.27E-04 | 2.57E-10 | 6.70E-07 | 1.20E-04 |

**Fig 5.**
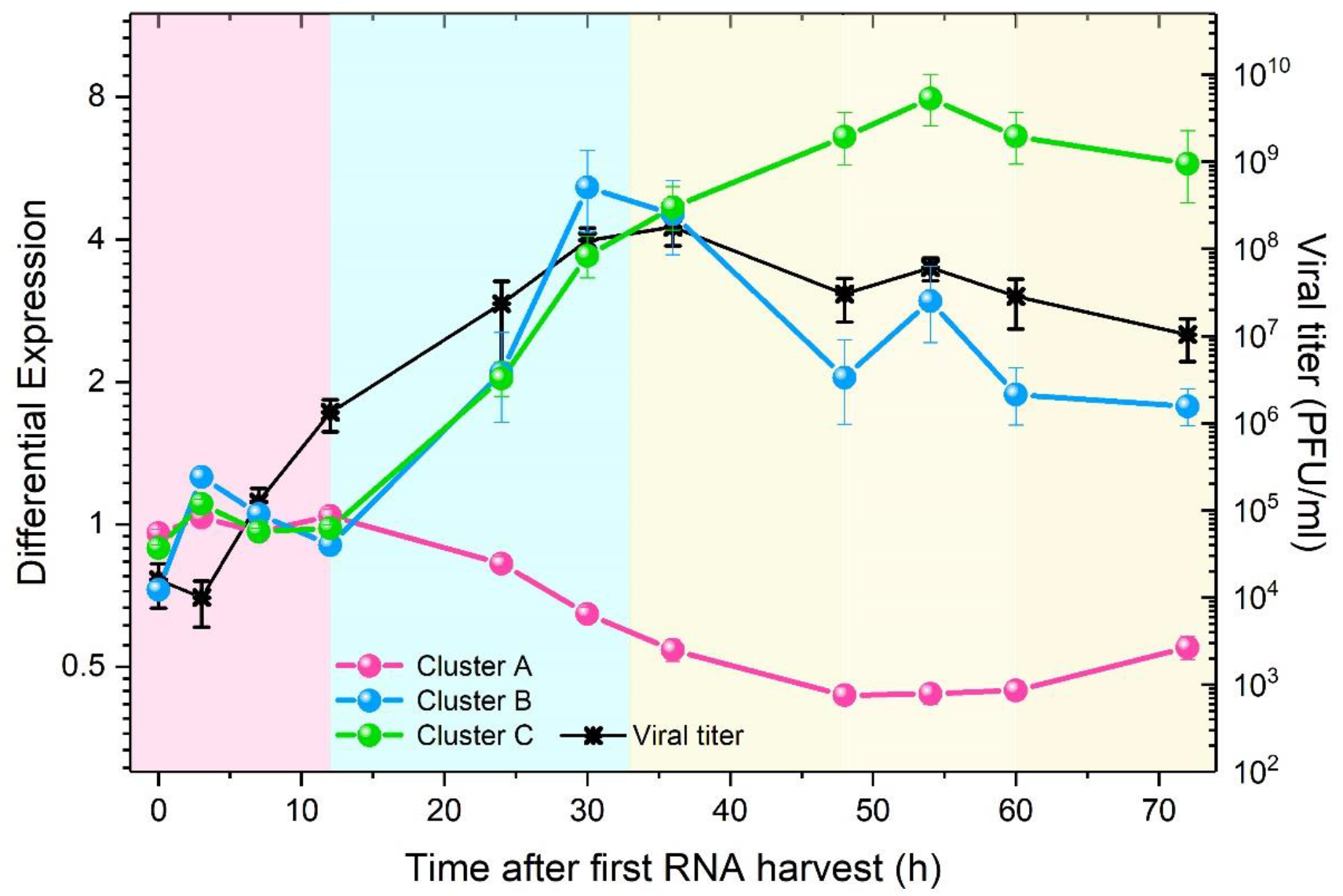
Time evolution of viral titer [14] (right log_10_ axis) and average differential expression of the covariant gene clusters A, B and C (left log_2_ axis). The control for each time point is the time-matched Mock sample. We identify three main phases for the host-virus interaction in the cell cultures. In the first phase, denoted by a pale-pink background, clusters A and C have differential expression near 1, while cluster B differential expression moves opposite to the viral titer. In the second phase, denoted by a pale-blue background, average differential expression in cluster A decreases monotonically while average differential expression in clusters B and C increases similarly, in parallel with increasing viral titer. In the third phase, denoted by a pale-yellow background, differential expression in clusters B and C diverges after the viral titer reaches its maximum, with Cluster B tracking viral titer evolution. After 54 h, however, viral titer and the average differential expression of clusters B and C decrease, while average differential expression in cluster A increases.

To characterize each cluster, we associate its genes with KEGG pathways and Gene Ontology: Biological Function terms using DAVID tools [19]. Additional files AF4A, AF4B, and AF4C in the Supplemental Information online present our full results. Cluster A (43 genes) is enriched with genes related to energy metabolism, *e*.*g*., Oxidation-Reduction and Retinol metabolism. As expected, this cluster shows reduced gene expression after viral infection. Cluster B (52 genes) is mainly enriched with terms related to cytokine production and the MAPK cascade. Finally, genes in Cluster C (124 genes), correlate with a broader spectrum of terms, but many genes relate to cytokine response (39 genes), signal transduction (62 genes) and regulation, including phosphorylation (60 genes). Terms or pathways can participate in more than one cluster simultaneously, but each gene participates in only one cluster. Genes in different clusters may, of course, interact.

Fig. 5 shows three distinct time courses for elements of the cells’ response to virus inoculation. The vertical axes represent the cluster average for gene differential expression in the *WT* samples *w*.*r*.*t*. the *Mock* samples. Genes in cluster B decrease (< 0 01) in expression relative to the control at *t* = 0 *h*. After 12 *h*, changes in differential expression increase rapidly, with 36 *h* and 54 *h* having (local) maxima in the collective differential expression dynamics. To test the significance of the difference of the fold-change we estimated the P-value for each time point using a two-tailed Welch test. We also calculated P values to assess significant differences in mean differential expression at each time point across clusters. We present these results in Table.

Fig. 5 shows the temporal pattern of response in the infected cells triggered by virus inoculation together with the time evolution of the viral titer, reproduced from Ref. [14], measured in units of PFU/ml (plaque forming units per milliliter) for 6 samples for each time point. We can observe that:

1. Viral titer initially decreases from 0 *h* to 3 *h*, then increases rapidly from 3 *h* to 36 *h*, decreases between 36 *h* and 48 *h*, then increases to a small, but statistically significant second maximum at 54 *h*, and finally decreases from 54 *h* to the end of the experiment.
2. In Cluster A (43 genes) average differential expression does not differ significantly from the control until 12 *h*. Between 12 *h* and 54 *h* average differential expression decreases, reaching a minimum at 54 *h*. After 54 *h*, average differential expression increases until the end of the experiment, but always remains less than 0.5. This cluster is enriched in genes involved in mitochondrial activity. Shi and collaborators showed that the SARS-CoV-1 protein designated *opening reading frame-9b* (ORF-9B) localizes to the outer mitochondrial membrane, manipulating host-cell mitochondria, and disturbing mitochondrial anti-viral signaling [20]. This interference could explain why Cluster A’s mean differential expression moves opposite to the viral titer.
3. The genes in Cluster B (52 genes) have the richest temporal dynamics of mean differential expression. At 0 *h*, differential expression is already depressed in the *WT* samples relative to the control, indicating that some genes change their expression very rapidly *w*.*r*.*t*. the control, during the 40 min incubation time before RNA harvesting (0 *h*). Average differential expression increases between 0 *h* and 7 *h*, then decreases until 12 *h*, then increases again, reaching a maximum at 30 *h*. Average differential expression then decreases until 48 *h*, then increases to a second, more modest maximum, at 54 *h* and finally decreases until the end of the experiment. Although the maximum at 54 *h* is modest in amplitude, it is statistically significant and coincides with the minimum value of the average differential gene expression in Cluster A and the second peak in viral titer, suggesting that it reflects a real change in biological function. The term enrichment analysis for Cluster B showed that 40 of the 52 genes in this cluster participate in Gene Ontology (*GO*) terms linked to “response to stimulus,” with 12 specifically tagged as “response to cytokines.” The great majority of the products of these genes localize either to the extracellular matrix, indicating signaling activity, or to the nucleus, indicating response to signaling. Other terms associated with genes in Cluster B link to ion transport.
4. In Cluster C (124 genes), after 12 *h* mean differential expression increases monotonically to a maximum at 54 *h*, after which it decreases until the end of the experiment. The maximum at 54 *h* coincides with the maximum at 54 *h* in Cluster B, the minimum in Cluster A, and the modest second peak in viral titer. Term enrichment analysis of the genes in Cluster C shows that the great majority of these genes have GO annotations involved in IFN response pathways, *e*.*g*., “I-kappaB kinase/Nf-kappaB signaling” (20 genes); other immune functions include “response to stimulus” (98 genes), “response to cytokine” (39 genes), “response to hormone” (19 genes) and “cytokine production” (31 genes).

Our analysis first considered Transcriptograms, which allowed us to select peaks in the differential Transcriptograms, representing 590 genes. That selection identified 219 genes with expression values (without Transcriptogram averages) that presented 2-fold or larger changes *w*.*r*.*t* time-matched controls at at least one point of the time series. This selection of relevant genes would not be possible without the previous use of the Transcriptograms. To show the difficulty of identifying biologically-relevant genes from the mass of statistically-significant genes using classical bioinformatics approaches, we applied two classical analysis approaches (Volcano plots and correlation with virus titer) and a more sophisticated one, Gene Set Enrichment Analysis (*GSEA*) [21], that searches the differentially-expressed gene sets based on a ranked gene list, ordered by differential expression as measured by a customized metric (with metrics including signal-to-noise ratio, P-value, and ratio-between-classes, amongst others). Supplementary Information online, sections 3 and 4, shows the results for classical bioinformatic analyses and Gene Set Enrichment Analyses (*GSEA*) comparing *WT* and *Mock* samples using three different combinations of data sets and metrics: signal-to-noise ratio and ratio-between-classes for normalized expression values and ratio-between-classes for values using Transcriptogram-averaged data. These Volcano plots, correlations and *GSEA* confirm the significance of the clusters found using our Transcriptogram analyses but, on their own, cannot identify these clusters directly from the original data, showing that the Transcriptogram analyses are more powerful in this context than the classical methods and *GSEA*. In the present analyses, Transcriptograms define a biological-relevance criterion, rather than a statistical-significance one, by suggesting the *scale* of differential expression at each time that should be considered for further investigation of statistical significance. Naturally, when increasing the threshold required to select gene sets to be further analyzed, and hence decrease the number of false-positive errors, we will simultaneously increase the incidence of false-negative errors. We stress that the present analysis tries to address problems caused by the abundance of false-positive errors in finding biologically-relevant genes in experiments showing genome-wide transcription alterations. The role played by this biological-relevance criterion in the selection of genes to be further analyzed becomes evident when we compare differential Transcriptograms at 0 *h* or 3 *h* with very modest peaks, to those at 30 *h* or later with much larger expression alterations (see Fig. 3 and Fig. S 7 in the Supplementary Information online). Although some peaks in the differential Transcriptograms may be statistically significant at early times, they are less relevant in comparison to the more pronounced peaks seen in differential Transcriptograms at later times.

In summary, we identified three gene clusters (A, B, and C) with distinct temporal profiles. Clusters B and C have the most distinctive patterns of temporal change, probably reflecting their specific functional roles during early infection. To verify the power of our method, we selected six genes from these clusters and analyzed their differential expression evolution. We then discuss their possible roles in the cellular response to virus inoculation. As we show below, these genes have well-known roles in immune response, showing that the Transcriptogram analysis was reasonable in identifying their variation as significant.

We begin by considering the gene pair HSD11B1 (from cluster C) and HSD11B2 (from cluster B). Both genes are linked to Cortisone-Cortisol balance. Cortisol is anti-inflammatory, secreted by the adrenal gland and present in plasma. It can be converted to inactive Cortisone by the enzyme 11-beta-hidroxysteroid dehydrogenase type 2 (HSD-2), the product of the HSD11B2 gene (For a review, see [21]). The time series for HSD11B2 in **Error! Reference source not found**. C (red line), shows that HSD11B2’s differential expression in infected cells begins to increase after 12 *h* up to 36 *h* reaching a 32-fold change *w*.*r*.*t*. the control, after which it gradually decreases to an 8-fold change. The reduction in anti-inflammatory Cortisol signaling between 12 *h* and 36 *h* probably enhances pro-inflammatory signaling in response to infection. HSD11B1 differential expression also starts increasing after 12 *h*, but peaks later, at about 54 *h*, as shown **Error! Reference source not found**. D (red line). 11-beta-hidroxysteroid dehydrogenase type 1 (HSD-1), the product of the gene HSD11B1, converts Cortisone back into Cortisol, possibly restoring anti-inflammatory signaling, after the initial pro-inflammatory response. The data thus show that the time evolution of these genes in this experiment reproduces the well-known interplay between Cortisone-Cortisol.

Fig. 4 D (olive green) highlights the differential expression evolution of JUN and TNFAIP3 from cluster C. JUN encodes c-Jun, a protein that participates in the transcription-factor complex “Activator Protein-1” (AP-1) that has complex context-dependent behaviors [22]. In epithelial cells, AP-1 components (containing c-Jun) may participate in regulating apoptosis or cell proliferation. JUN differential expression seems to increase with viral titer after a few hours of delay. Cell-cycle mean expression is depressed in both *Mock* and *WT* samples compared to *Mock* sample expression at 0 *h* (the interval marked A2 in Fig. 2 and Fig. S 6 in the supplemental information online), probably due to contact inhibition in the culture. However, Fig. S 6 shows that cell-cycle gene expression recovers after 60 *h*. Both apoptosis and proliferation may occur in the infected culture. The observed time series for JUN differential expression may relate to these differences in cell cycle-related expression between *WT* and *Mock* samples.

TNFAIP3 encodes the protein A20, a negative regulator of the NF-κB protein complex. TNFAIP3 is thus a negative regulator of inflammation and is known to be rapidly induced after Toll-like receptors interact with a pathogen or respond to TNF-α or IL-1 cytokines [23]. **Error! Reference source not found**. D shows a peak of differential expression of TNFAIP3 at 48 *h*, followed by monotonic decrease until the end of the experiment. Comparing the time-series for the differential expression of TNFAIP3 shown **Error! Reference source not found**. 4 D with the viral titer evolution in Fig. 5 5, we find that TNFAIP3 expression follows the viral titer after about a 4 *h* delay. This temporal relationship suggests that the anti-inflammatory response due to TNFAIP3 in the *WT* sample gradually decreases as the viral titer decreases.

Fig.4 C shows the dynamics of differential expression of individual genes in cluster B. We have highlighted in navy blue the expression of EGR3 and TWIST1, the two genes whose differential expression presents the largest fold changes in cluster B. TWIST1 negatively regulates the NF-κB protein complex. TWIST1 is thus anti-inflammatory [24]. The variation in the TWIST1 time series in **Error! Reference source not found**. C generally follows the viral titer evolution in Fig. 5. EGR3 is a zinc-finger transcription factor of the Early Growth Transcription family (EGR) that responds early to environmental stimuli to induce cell proliferation, differentiation, and immune responses [25]. In resting epithelial cells, EGR3 is usually weakly expressed, but a wide variety of extracellular signals such as cytokines and T-cell receptor (TCR) activation can promote EGR3 expression [25]. Fig 4 C shows that EGR3 differential expression increases after 12 *h* and remains high, varying between 16- and 32-fold change from 20 *h* to 54 *h*, then decreasing when the viral titer begins to decrease after 54 *h* (Fig. 5). This correspondence suggests that the virus may activate EGR3 in epithelial cells. Hypothesizing that the virus also promotes EGR3 in T cells, fibroblasts, and endothelial cells could explain T-cell anergy in SARS-CoV-1 and SARS-CoV-2 infection in T cells, since the co-activation of T-cell receptors by antigen and EGR3 may lead to T cell anergy [27,28]. Furthermore, EGR3 regulates fibrogenic responses in fibroblasts [29], and EGR3 may cause vascular disruption when active in vascular endothelial cells [29]. The infected lung contains epithelial cells, fibroblasts and endothelial cells, all of which express the ACE-2 receptor for SARS-CoV-1 and SARS-CoV-2 [30,31,32,33]. T-cells have also been reported as being infected in SARS-CoV-2 in a preprint in biorXiv [34]. Both T-cell depletion due to exhaustion or anergy, and fibrotic sequels have been reported in SARS-CoV-1 [35] and SARS-CoV-2 [36] patients. We wonder whether these effects on T-cells and fibroblasts may correlate with the activation of EGR3 by the virus. Also, since EGR3 activates VEGF in endothelial cells [29], its activation in infected cells may link to the endothelialitis, thrombosis, and angiogenesis reported in COVID-19 [5]. We stress, however, that the six highlighted genes are only examples that deserve further examination. The other genes in clusters A, B, and C also deserve further exploration.

Finally, in Supplementary Information online, Figs. S 8 and S 9 present the Transcriptograms for the *DORF6* sample data sets produced by Sims *et al*. in the same experiment which generated the *WT* and *Mock* data series. Fig. S 10 shows the evolution of the mean expression in clusters A, B, and C for *DORF6* samples, showing that genes in cluster B carry the main differences between the immune response to *DORF6* and *WT* infection.

## Conclusions and Perspectives

Transcriptogram analysis of microarray time series experiments by Sims *et al*. [14] for SARS-CoV-1 infection of Calu3-2B4 cells, a human epithelial cell line selected for ACE-2 expression [8], identifies three main gene sets with well-defined dynamics, summarized in Fig.5. Differential expression profiles indicate that the average differential expression for Clusters A, B, and C are more intense from 12 *h* after inoculation, and that mitochondrial activity (ClusterA) decreases until 54 *h*, then partially recovers. Clusters B and C both consist mostly of genes associated with immune response, and their averages show a marked increase beginning at 12h after inoculation. However, their dynamics in response to viral inoculation differs, and while Cluster B presents an average that decreases after 30h, following virus titer, cluster C average keeps increasing up to 54h post infection. Cluster B consists mainly of genes related to innate immune response. Cluster C comprises genes related to both innate and adaptive immune responses. Beyond pro/anti-inflammatory signaling via, for example, the negative regulation of NF-κB complex by TNFAIP3 and the interplay between HSD11B1 and HSDB11B2 differential expression, the identity of the genes in each cluster suggests that the response to viral inoculation also includes regulation of apoptosis and proliferation, via JUN, and has secondary effects on cell differentiation (with different possible outcomes, depending on the cell type), via EGR3. These effects follow the temporal patterns of either Cluster B or Cluster C, suggesting coordinated cellular responses. Because the Transcriptogram analysis selects genes most functionally relevant to the specific behavioral changes in a particular experiment, we could identify the correlated responses of genes associated with different pathways or GO terms, which would be hard to identify if we conducted correlation analyses of the temporal expression changes of all genes at once. The differential Transcriptogram, by identifying differential expression bands of functionally related genes, greatly reduces the number of “genes of interest” making their detailed temporal analysis practical. Because mathematical models usually consider variables that aggregate the effect of multiple genes into broad representations of classes of biological mechanisms or pathways, these mean differential expression time series can serve as direct validation data for mathematical models of epithelial-cell responses to SARS-CoV-1 infection.

Because gene expression changes in control samples in cell culture as well as in infected samples, using time-matched gene expression controls is critical to distinguish cell-culture effects from infection effects.

EGR3 activation may explain several symptoms in patients with severe responses to SARS-CoV-1 and SARS-CoV-2 infection. Other cell types besides epithelial cells may be infected by the virus in the infected lung: in particular, if EGR3 activation also occurs in infected T-cells, it could explain T-cell anergy (against viral antigens), in infected fibroblasts, EGR3 activation could link to observed fibrosis, and in infected endothelial cells, if EGR3 activation could explain the endothelialitis, thrombosis, and angiogenesis reported in COVID-19. We suggest that these hypotheses would be worth investigating in future experiments.

Our analysis identified as differentially expressed two genes from a well-known feed-back loop which regulates Cortisol-Cortisone balance. Infection perturbs the resting Cortisone-Cortisol homeostasis and we would expect that each gene would follow a different time course in response to infection. We find that differential expression of the proinflammatory HSD11B2 follows Cluster B and peaks with viral titer, while the anti-inflammatory HSD11B1 follows Cluster C and peaks later. We will examine the remaining 213 genes identified as significant by our Transcriptogram analsysis in future work

Transcriptograms allowed us to define an appropriate context-dependent threshold for mechanistic relevance of gene differential expression. If we knew, *a priori*, which genes are relevant, both classical and *GSEA* analyses would yield the same results as our Transcriptogram analysis. However, without knowing in advance which genes are relevant, classical analyses detect <u>every</u> gene with statistically-significant differential expression, leaving us with too many genes and hypotheses to be useful. The previous filtering using Transcriptograms reduced the number of genes of interest to a tractable set, allowing us to suggest shared mechanistic functions for the observed gene expression patterns. These gene sets are defined by the data directly, not by reference to previously defined pathways or biological functions.

Transcriptogram analysis selects and aggregates the biologically-relevant components of the experimental time series in a way that will support detailed mathematical modeling of the inflammatory response of lung cells to SARS-CoV. This approach to aggregating time-series information to yield quantitative net functional responses is new and useful for the complex undertaking of explaining virus effects on inflammatory response, an explicit goal of, for example, the NIH new IMAG/MSM Working Group for multiscale in-host modeling of viral infection and immune response (https://www.imagwiki.nibib.nih.gov/working-groups/multiscale-modeling-and-viral-pandemics).

We could apply the same methodology to identify functional differences between cell-culture responses to SARS-CoV-1 infection between male-derived and female-derived cells or between adult-derived and juvenile-derived cells (to identify sex-linked and age-linked changes in response pattern). The same methods could identify critical differences in cell responses to SARS-CoV-1 and SARS-CoV-2 infection or among responses to infection by other respiratory viruses. We could also study differences in response between cells derived from different possible loci of infection (nasal, throat, bronchial, alveolar, heart, kidney), or to compare infection responses between classical cell culture and organoids, between organoids derived from different donors, or between different initial infection intensities.

## Acknowledgement

This work has received support from Brazilian agencies CNPq and CAPES. JAG acknowledges support from the Falk Medical Research Trust Catalyst Program and the US National Institutes of Health, grants U01 GM111243, R01 GM076692 and R01 GM077138.

## Authors Contributions

RdA and GLT ran the analyses. RdA, GLT, and JAG designed and wrote the paper.

## Corresponding Author

Rita de Almeida.

## Competing interests

JAG discloses that he owns shares in Gilead Sciences Inc. GLT and RdA declare no potential conflict of interest.

## Code and Data availability

The analyses have been performed on a previously published dataset by Sims and collaborators [14]. The dataset is publicly available Gene Expression Omnibus (*GEO*) under accession number GSE33267 (https://www.ncbi.nlm.nih.gov/geo)

The program we used for our analyses is freely available at https://lief.if.ufrgs.br/pub/biosoftwares/transcriptogramer/ and is also available as an R-Bioconductor function. The ordered gene list we used is available as a text file and is included in the Supplementary Information online.

## Figure Legends

**Fig. 6.** *Gene list and enrichment of terms related to critical biological functions as a function of position in the list. From left to right, in shades of purple, the list is enriched with genes associated with translation and mRNA processing then pathways linked to the cell cycle. Next, in shades of blue, genes associated with cell differentiation and, in shades of green, genes associated with immune response, cytokine production and interaction with the extra-cellular matrix (ECM). Finally, shades of orange denote genes associated with energy metabolism.*

**Fig. 7.** *Relative Transcriptograms of radius 30 for Mock and WT samples, using the Mock sample’s expression at t = 0 has the control. The labeled time is the experimental time after the first RNA harvest (over 40 minutes after inoculation). Vertical axes are on a log_2_ scale. Green horizontal lines in the bottom panel correspond to 9/5 and 5/9 fold changes. Black horizontal lines represent the control sample (Mock) expression. Red and blue lines represent the relative Transcriptograms for, respectively, WT and Mock samples. Gray, light red and cyan shading indicate the standard errors of the respective relative Transcriptograms. We identify 11 intervals, indicated by the horizontal color bars, where the red line differs from the control by more than 9/5 at 48 hor after, except for peak A1. (See the panel for the complete time series in Supplementary Information online, Fig. S 6).*

**Fig. 8.** *Differential Transcriptograms WT(t)/Mock(t) (radius 30). Time is time after the first RNA harvest. Vertical axes are on a log_2_ scale. Black horizontal lines represent the control sample Mock(t)/Mock(t). Violet lines are the differential Transcriptograms for WT(t)/Mock(t). Light violet shading indicates standard errors for WT Transcriptograms. We identify 17 bands where the violet line differs from the control more than 9/5-fold at 24 h. The horizontal red lines denote the 9/5-fold and 5/9-fold lines in all panels.*

**Fig. 9.** *Covariance matrix for the time changes of differential expression (time-matched control) for 219 genes selected from the significantly differentially expressed Transcriptogram bands identified in Fig. 3. We find 3 major covariant clusters. Table 1 lists the genes in each cluster. Figs. 4 B, C, and D show the time-series for the differential expression for each gene in each cluster and the averaged differential expression time-series for each cluster. Selected genes in each cluster are highlighted. The control for each gene at each time point is the Mock sample expression of the same gene at a matched time. Expression is for individual genes, not Transcriptogram averages over the neighbors in the ordered gene list. Clusters A (43 genes), B (52 genes), and C (124 genes) have highly covariant differential expression time series. Additional file AF2, in the Supplementary Materials online shows the individual relative expression time series for each gene in full detail.*

**Fig. 10.** *Time evolution of viral titer [14] (right log_10_ axis) and average differential expression of the covariant gene clusters A, B and C (left log_2_ axis). The control for each time point is the time-matched Mock sample. We identify three main phases for the host-virus interaction in the cell cultures. In the first phase, denoted by a pale-pink background, clusters A and C have differential expression near 1, while cluster B differential expression moves opposite to the viral titer. In the second phase, denoted by a pale-blue background, average differential expression in cluster A decreases monotonically while average differential expression in clusters B and C increases similarly, in parallel with increasing viral titer. In the third phase, denoted by a pale-yellow background, differential expression in clusters B and C diverges after the viral titer reaches its maximum, with Cluster B tracking viral titer evolution. After 54 h, however, viral titer and the average differential expression of clusters B and C decrease, while average differential expression in cluster A increases.*

## Notes

### Competing Interest Statement

The authors have declared no competing interest.

### Summary of Updates

We have revised the first version to include the referees' suggestions.

