## Supplementary information for "Transcriptogram analysis reveals relationship between viral titer and gene sets responses during Corona-virus infection"

### 1 Experiment and data description (after Reference [1])

Sims *et al*. [1], made their raw microarray data accessible through the Gene Expression Omnibus (GEO) web site under accession number GSE33267. Their cell lines consisted of clonal population of Calu3 cells (human lung adenocarcinoma cells) sorted for high levels of expression of ACE2, (angiotensin-converting enzyme 2), which they referred to as Calu3 2B4 cells. They grew the Calu3 2B4 cells in minimal essential media (MEM; Invitrogen-Gibco) containing 20% fetal bovine serum (HyClone) and 1% antibiotic-antimycotic mixture (Invitrogen-Gibco). They performed viral titration assays in Vero E6 cells, maintained in MEM (Invitrogen-Gibco) containing 10% fetal clone II (HyClone) and 1% antibiotic-antimycotic (Invitrogen-Gibco). They derived their icSARS-CoV virus (both *WT* and *DORF6* strains) from Prof. R. Baric´s laboratory infectious clone constructs as described in Refs. [1,2,3,4].

They plated Calu3 2B4 cells in triplicate, washed them prior to infection, and either infected them at a multiplicity of infection of 5 (MOI 5) (*WT* and *DORF6* samples) or inoculated them with *Mock* solutions (*Mock* samples), then incubated them at 37°C for 40 min. They then removed the inoculum, washed cells 3 times with 1× phosphate-buffered saline (PBS), and added fresh medium prior to their defined time zero. The time points for these procedures were $0, 3, 7, 12, 24, 30, 36, 48, 54, 60,$ and $72$ $h$ post-infection They collected medium from each well to determine viral titers at each time point. For microarray analysis, they washed the cells in 1 × PBS, then harvested them in TRIzol (Invitrogen) and stored them at -80°C.

For the microarray analyses, all probes were required to pass Agilent quality control (QC) flags for all replicates of at least one infection time point (33,631 probes passed).

#### 1.1 Original results comparing *WT* SARS-CoV infected and *Mock* samples (after Reference [1])

Viral titers increased by 4 orders of magnitude by $30$ $h$ post-infection in *WT*-samples, with peak titers reaching $10^8$ pfu/ml at $36$ $h$ post-infection.

The original paper [1] determined differential expression by comparing icSARS-CoV-infected replicates to *Mock*-(un)infected replicates for each time point, based on a fit to a linear model for each transcript. They set the criterion for differential expression as an absolute $\log$ 2-fold change of $1.5$ and a false discovery rate (FDR)-adjusted *P* value of $0.05$ for each time point.

They harvested total RNA in triplicate at the time points described above, and analyzed global mRNA expression. They detected little cellular gene differential expression in comparison with *Mock* samples during the first 24 $h$ of icSARS-CoV infection.

### 2 Normalization check

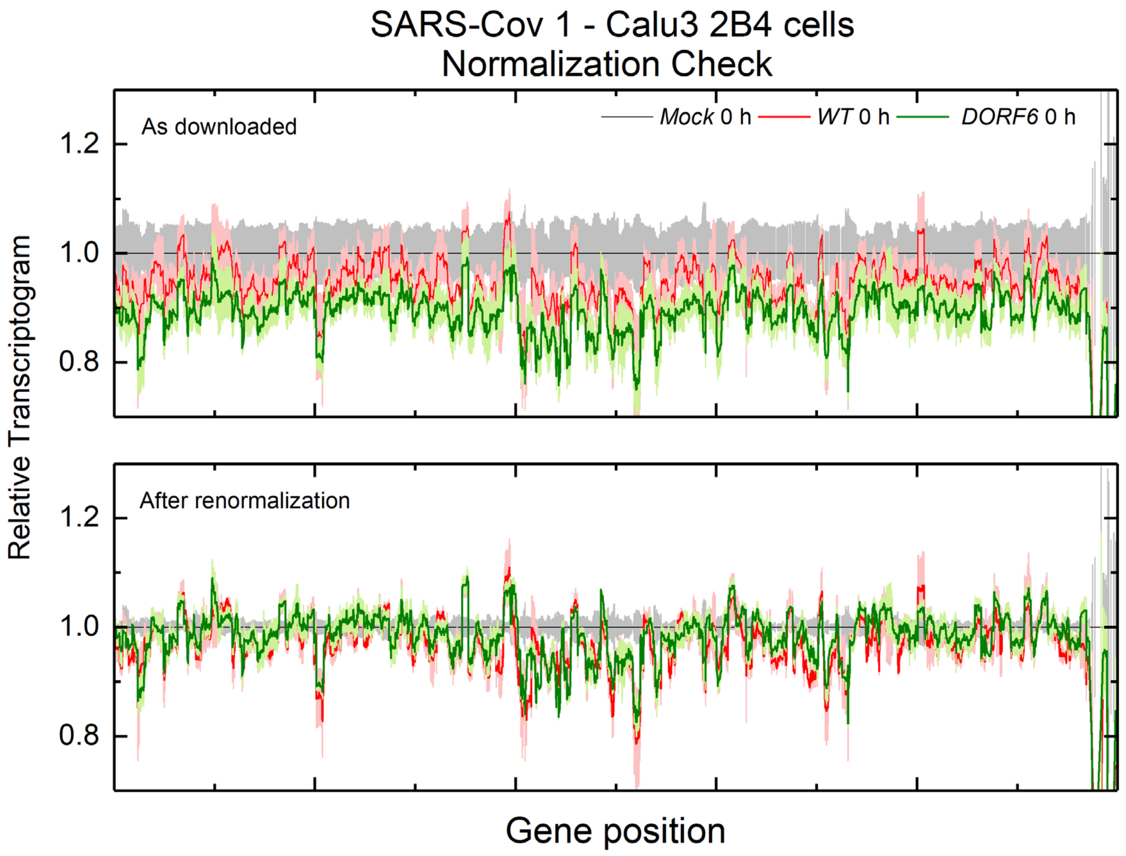


*Figure S 1. Upper panel: Relative Transcriptograms (Radius 30) using Mock $0\ h$ as a control for both WT at $0\ h$ and DORF6 (mutant strain that does not express ORF6 protein) at $0\ h$ for expression data downloaded from the Gene Expression Omnibus site. The offset in the raw data is clear. Bottom panel: Transcriptograms produced from expression data re-normalized so that the average expression for each sample is one, eliminating the offset. All analyses in this paper used re-normalized data.*

### 3 Classical analysis

#### 3.1 Volcano plots

We produced Volcano plots comparing time-matched Wild Type samples against *Mock* samples for all 11 time points. As expected, the locations in the plots of the genes in clusters A, B, and C change with the time after first RNA harvest. Before 24 $h$, most genes in these clusters are not differentially expressed. At and after 24 $h$, most genes in cluster A and most genes in cluster B localize to the down-regulated branch (left branch), while genes in cluster C mostly localize to the up-regulated branch (right branch). Genes in cluster B follow the trend of cluster C before 48 $h$, after which they localize in regions with more modest values of fold change. The localization of the genes from different Transcriptogram-identified clusters to different compact regions in the Volcano plots supports the validity of our Transcriptogram clustering. However, without the Transcriptogram analysis, we could not have identified these genes as functionally significant, because they would be hidden among the many other genes that the Volcano plots find are differentially expressed and assign to the same regions.

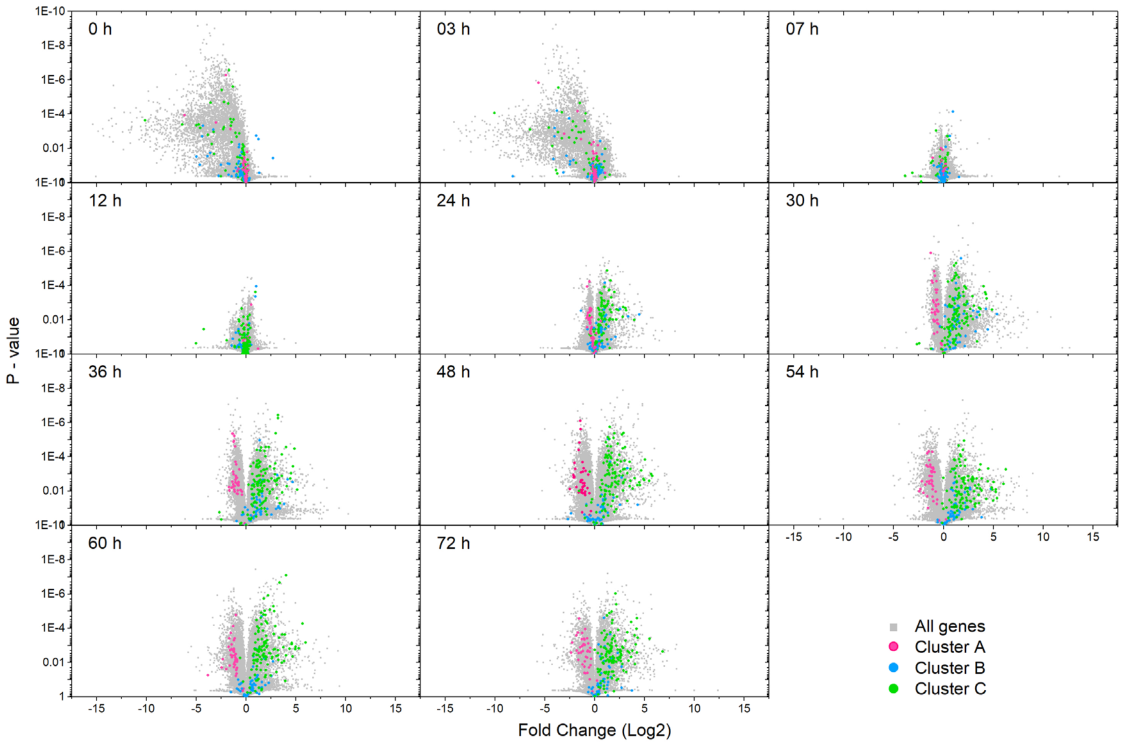


*Figure S 2. Volcano plots (P-value versus Fold-Change) for all genes with probes in the microarrays, comparing time-matched WT samples to Mock samples. We highlight the genes belonging to the clusters found with the Transcriptogram method in pink (Cluster A), blue (Cluster B) and green (Cluster C). The Transcriptogram clusters populate the regions with low P-values and large fold changes in the Volcano plots, supporting the statistical significance of their differential expression. However, the Volcano plots also find many other genes which overlap these clusters, which the Transcriptogram analysis does not find to be biologically relevant.*

#### 3.2 Correlation between gene expression and virus titer

For all genes with probes in the microarray, we calculated the time correlation $CVirus_i$ between the differential expression of the $i$-th gene and the virus titer, that is:

$$CVirus_i = \frac{1}{11}\sum_{t=1}^{11}\frac{[e_i(t)-\overline{e_i}]\times[\tau(t)-\overline{\tau}]}{\sigma_i\sigma_{virus}},$$

where $e_i(t)$ is the differential expression defined in Eq.(1) in the main text (the ratio between the average expression of *WT* samples and *Mock* samples at time *t*), $\overline{e_i}$ is the time-average of differential expression for the $i$-th gene, and $\sigma_i$ is the corresponding standard deviation. $\tau(t)$ is the virus titer at time $t$, $\overline{\tau}$ is the virus titer time average and $\sigma_{virus}$ the corresponding standard deviation. Figure S 3 shows $CVirus_i$ for all genes, in random order. We have also highlighted the genes belonging to clusters A (pink), B (blue), and C (green). Not surprisingly, genes in cluster B have the highest correlation with virus titer: their differential expression increases and decrease in parallel with changes in the virus titer. Differential expression of genes in cluster C usually has positive correlation with changes in viral titer, but the correlation is weaker than for genes in cluster B, since their differential expression keeps increasing after the virus titer starts to decrease. Finally, differential expression of genes belonging to cluster A correlates negatively with changes in virus titer. These results agree with those found using the Transcriptogram method, but without the Transcriptogram analysis, we could not have identified these functionally relevant genes among the host of other genes whose differential expression has similar net time correlations with changing virus titer.

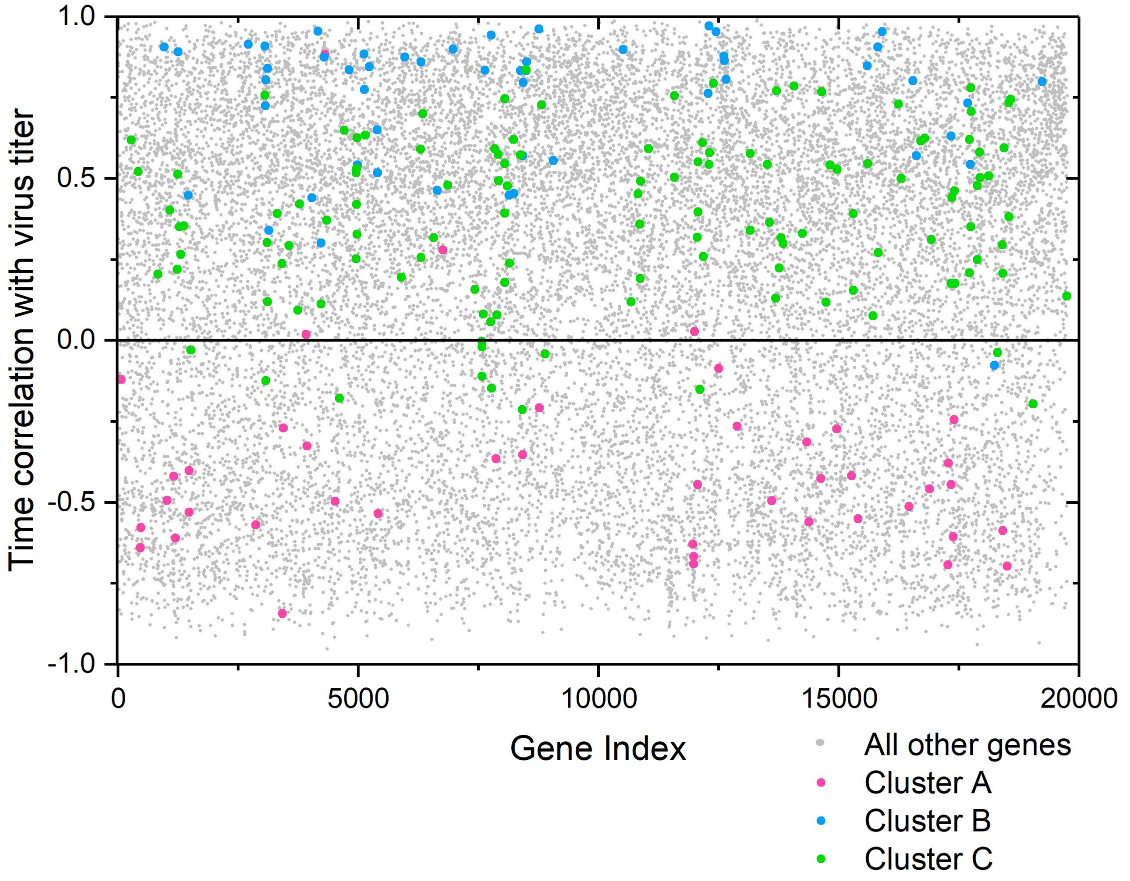


*Figure S 3. Time correlation between differential expression and virus titer for all genes with probes in the microarray. We highlight the genes belonging to the clusters found using the Transcriptogram method in pink (Cluster A), blue (Cluster B) and green (Cluster C).*

Figure S 4 superimposes the time correlation between differential expression and virus titer on the Volcano plots: the Volcano-plot points are colored by their correlation value. At early times, genes that correlate positively and negatively with virus titer are mixed together in the Volcano plot. Positively- and negatively-correlated genes segregate after $24\ h$, with maximal separation at $30\ h$ and $36\ h$, with positively-correlated (red) genes lying in the up-regulated branch of the Volcano plots and negatively-correlated (blue) genes lying in the down-regulated branch. At later times, some positively-correlated genes migrate to the down-regulated region of the Volcano plots though the overall separation is maintained. Figure S 4 again shows that classical analysis, by itself, cannot identify biologically meaningful genes.

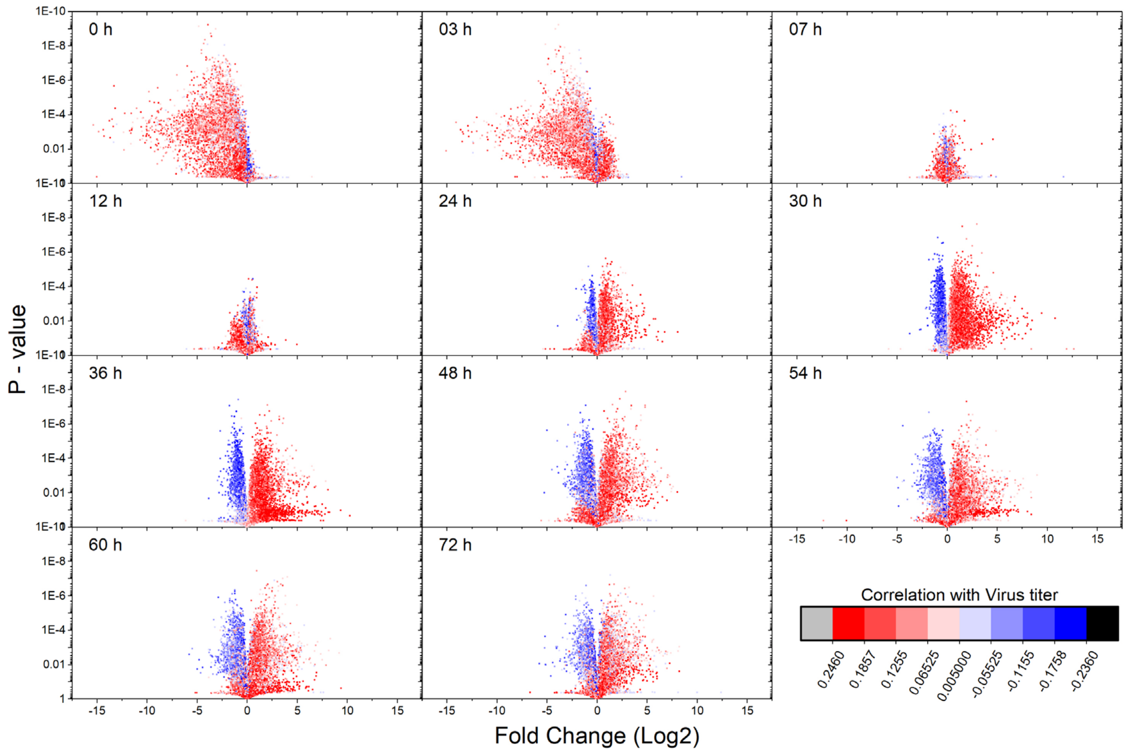


*Figure S 4. Volcano plots (P-value versus Fold-Change) for all genes with probes in the microarrays, comparing time-matched WT samples to Mock samples. We have colored each points by the gene's time correlation between its differential expression and the virus titer, following the color bar in the right lower corner of the panel. Before 12 $h$, genes with positive (red) and negative (blue) temporal correlation with the viral titer occur in both the up-regulated and down-regulated branches of the Volcano plots. At 30 $h$ and 36 $h$ the positively-correlated genes mostly lie in the up-regulated branch, while the negatively-correlated genes lie in the down-regulated branch. Overall segregation persists but gradually weakens at later times.*

#### 3.3 t-test for differential gene expression in the clusters

Here we present t-tests for the mean expression of the clusters, comparing Wild Type samples to *Mock* samples. Although the Transcriptograms suggest that mean differential expression (as defined in Eq. (1) in the main text) of these clusters before 12 $h$ are not relevant, and genes with positive and negative temporal correlation with virus titer are present in both branches of the Volcano plots, t-tests indicate that differential expression of genes in cluster B, calculated between the Wild Type and *Mock* samples is statistically significant, presenting a low P-value (see Table S1, below). However, while statistically significant, this differential expression is probably not functionally relevant in this experiment. The t-test is overly sensitive and is detecting small variations that are not that significant to the cells' immune response to viral infection. After 24 $h$, the differential expression between the clusters is both statistically significant and biologically relevant, as indicated by our Transcriptogram analyses and by the segregation of genes positively and negatively correlated with the virus titer in the Volcano plots.

| Time (h) | Cluster A | | | | |
|---|---|---|---|---|---|
| | Mock Samples | | WT Samples | | |
| | Mean | Standard deviation | Mean | Standard deviation | P value |
| 0 | 0.002106 | 2.79E-05 | 0.001916 | 8.87E-05 | 0.054983 |
| 3 | 0.002072 | 9.17E-06 | 0.001828 | 2.67E-05 | 0.00178 |
| 7 | 0.001901 | 6.31E-05 | 0.001834 | 4.37E-05 | 0.215733 |
| 12 | 0.001915 | 1.00E-05 | 0.001914 | 4.79E-05 | 0.963999 |
| 24 | 0.002128 | 0.000103 | 0.001557 | 1.58E-05 | 0.009353 |
| 30 | 0.002152 | 1.31E-05 | 0.001198 | 1.96E-05 | 0.022868 |
| 36 | 0.002224 | 3.09E-05 | 0.001029 | 4.62E-05 | 1.15E-05 |
| 48 | 0.002198 | 6.90E-05 | 0.000925 | 3.14E-05 | 0.000148 |
| 54 | 0.002202 | 0.000126 | 0.000876 | 5.50E-05 | 0.000784 |
| 60 | 0.002215 | 0.000162 | 0.000924 | 4.49E-05 | 0.003143 |
| 72 | 0.002176 | 7.62E-05 | 0.001175 | 4.63E-05 | 0.000162 |

| Time (h) | Cluster B | | | | |
|---|---|---|---|---|---|
| | Mock Samples | | WT Samples | | |
| | Mean | Standard deviation | Mean | Standard deviation | P value |
| 0 | 0.000403 | 2.32E-05 | 0.000147 | 1.56E-05 | 0.000219 |
| 3 | 9.15E-05 | 9.39E-06 | 4.11E-05 | 3.74E-06 | 0.00541 |
| 7 | 5.64E-05 | 2.41E-06 | 8.51E-05 | 1.27E-06 | 0.000335 |
| 12 | 4.87E-05 | 2.54E-07 | 7.23E-05 | 7.92E-06 | 0.035293 |
| 24 | 5.08E-05 | 4.69E-06 | 0.000305 | 2.95E-05 | 0.003713 |
| 30 | 4.46E-05 | 3.87E-06 | 0.000548 | 0.000134 | 0.001307 |
| 36 | 5.20E-05 | 2.71E-06 | 0.000564 | 6.08E-05 | 0.004604 |
| 48 | 5.96E-05 | 2.68E-06 | 0.000345 | 1.17E-05 | 0.000311 |
| 54 | 5.43E-05 | 4.19E-06 | 0.000251 | 8.71E-06 | 6.93E-05 |
| 60 | 8.19E-05 | 2.74E-06 | 0.000215 | 2.23E-05 | 0.008514 |
| 72 | 6.55E-05 | 7.56E-06 | 0.000126 | 3.63E-06 | 0.001338 |

| Time (h) | Cluster C | | | | |
|---|---|---|---|---|---|
| | Mock Samples | | WT Samples | | |
| | Mean | Standard deviation | Mean | Standard deviation | P value |
| 0 | 0.003577 | 5.82E-05 | 0.002238 | 5.17E-05 | 8.59E-06 |
| 3 | 0.002784 | 6.95E-05 | 0.001822 | 8.86E-05 | 0.000173 |
| 7 | 0.002001 | 8.67E-05 | 0.002016 | 6.55E-05 | 0.825886 |
| 12 | 0.001842 | 4.54E-05 | 0.001891 | 5.27E-05 | 0.289381 |
| 24 | 0.001802 | 0.000107 | 0.003337 | 0.000169 | 0.000489 |
| 30 | 0.001776 | 5.02E-05 | 0.004903 | 0.000238 | 0.001307 |
| 36 | 0.001877 | 9.05E-05 | 0.006397 | 0.000297 | 0.000617 |
| 48 | 0.001896 | 2.87E-05 | 0.009491 | 6.60E-05 | 1.15E-06 |
| 54 | 0.001869 | 0.000115 | 0.009533 | 0.000334 | 0.000183 |
| 60 | 0.001922 | 0.000131 | 0.009321 | 0.000305 | 8.43E-05 |
| 72 | 0.001868 | 2.90E-05 | 0.007499 | 0.000144 | 0.000127 |

*Table S 1. Average Expression (over samples and genes) for genes in clusters A, B and C for the 11 time points for both Mock and WT samples. The difference in expression between Mock and WT genes in cluster B Mock is significant as early as $0\ h$ after the first RNA harvest.*

### 4. GSEA and Transcriptogram results

The classical analyses confirm the results obtained using Transcriptograms, but would not, on their own be sufficient to produce gene sets A, B and C. Here we present results using a bioinformatic tool specially designed for gene-set analysis, Gene Set Enrichment Analysis (*GSEA*) [5].

Gene Set Enrichment Analyses (*GSEA*) investigate differential expression in genome-wide microarray or RNA-Seq transcription data. After quality control and normalization, *GSEA* can test for differential mean expression of gene sets between two classes of samples. Gene sets may be custom-built or use public data bases, such as Gene Ontology terms or KEGG pathways. *GSEA* first produces a gene list, ranking each gene by one of several possible metrics used to express the differential expression between genes of two classes (*WT* versus *Mock*, for example). The default metric is *Signal-to-Noise ratio* (calculated as the difference between each gene expression averaged over the replicates of each class divided by the sum of the corresponding standard deviations) but “Ratio between Classes” (calculated as the ratio between the gene expression average over replicates for each class) is also an option. The second step locates all genes from a given gene set (for example, the sets of genes belonging to a KEGG metabolic pathway or a term in the Gene Ontology) within the ranked list. If genes from a gene set are predominantly located in high- (low-) scoring regions of the ranked gene list, the gene set is identified as differentially up(down)-expressed. The significance of the identification may be estimated by calculating the False Discovery Rate (*FDR*) calculated under permutation of either the samples or the genes. For sample permutation, identification is accepted as significant when the FDR < 25%. For gene-list permutation, identification is accepted as significant when the FDR < 5%. When all classes have few replicates (less than 7), the acceptance criterion is based on gene-set permutation.

We ran *GSEA* analyses for the 11 time points comparing *WT* and *Mock* classes, with the following selection of metrics:

a. Expression data, Signal-to-Noise metric, gene-set permutation, testing all GO:Biological Function terms.
   Expression data, Ratio-between-Classes metric, gene-set permutation, testing all GO:Biological Function terms.

Like *GSEA*, Transcriptograms help identify biologically-meaningful genes in the presence of noisy signals. We applied *GSEA* to the gene sets identified by the Transcriptograms of radius 30 (as in the original analysis). This workflow considers only genes that are in the Transcriptogram ordered gene list. The Transcriptogram’s running-window average reduces signal noise and assign a value to each gene in the ordering. We then ran *GSEA* for the Transcriptogram values as follows:

b. Transcriptogram values (after averaging over neighbors) with radius 30, Ratio-between-Classes metric, gene permutation, for all GO:Biological Function terms.

Table S 2 summarizes the results of the *GSEA* and Transcriptogram/*GSEA* analyses.

| Number of GO:BP terms identified as significant at FDR < 5% | | | | | | |
|---|---|---|---|---|---|---|
| Time after first RNA harvest | Expression data: Signal-to-Noise | | Expression data: Ratio-between-Classes | | Transcriptogram-data: Ratio-between-Classes | |
| | *WT* up | *Mock* up | *WT* up | *Mock* up | *WT* up | *Mock* up |
| $0\ h$ | 44 | 12 | 4 | 13 | 61 | 255 |
| $3\ h$ | 0 | 33 | 0 | 409 | 32 | 41 |
| $7\ h$ | 0 | 0 | 0 | 7 | 28 | 298 |
| $12\ h$ | 7 | 2 | 0 | 16 | 73 | 67 |
| $24\ h$ | 266 | 133 | 0 | 126 | 769 | 184 |
| $30\ h$ | 205 | 179 | 0 | 120 | 942 | 161 |
| $36\ h$ | 461 | 238 | 0 | 72 | 973 | 165 |
| $48\ h$ | 799 | 129 | 1 | 21 | 977 | 150 |
| $54\ h$ | 685 | 99 | 0 | 22 | 920 | 152 |
| $60\ h$ | 856 | 66 | 0 | 17 | 871 | 157 |
| $72\ h$ | 718 | 13 | 0 | 16 | 388 | 98 |

*Table S 2. Number of terms from Gene Ontology: Biologic Function found differentially expressed with significance at FDR < 5% by GSEA, up-regulated in either the WT class (WT up) or in Mock class (Mock up).*

*GSEA* results depend dramatically on the choice of metric and data pre-processing. Comparing the two metrics applied to the normalized data, at $24\ h$ after first RNA harvest and after, *GSEA* with the Signal-to-Noise metric finds hundreds of significantly differentially-expressed terms, while *GSEA* with the Ratio-between-Classes metric finds essentially none. This sensitivity to the metric used is a concern for the reliability of the method. Using the Transcriptogram to smooth the data with a running window average over the ordered gene list decreases noise. Consequently, the number of terms found to be significantly differentially-expressed increases. In fact, the *ES* (enrichment score) for each gene produced by *GSEA* applying Ratio-between-Classes to Transcriptogram-smoothed data is *identical* to the relative Transcriptogram presented in Figure 3 in the main text or Figure S 6 below, differing only in the order of genes: the assigned value for each gene is the same. The hundreds of terms from GO:BP which *GSEA* finds to be differentially expressed are *statistically significant* in the sense that their difference in expression is much larger than their standard deviation. However, we may expect that many of these statistically-significant terms are not *relevant* to the biology of the problem (for example, 409 terms were found up-regulated in *Mock* samples, when applying GSEA to expression data at $3\ h$ after RNA harvest). The Transcriptograms in Figure S 6 show the variation in scale of the differential expression of different gene sets: the alterations at, for example, $7\ h$ after first RNA harvest are small compared to those observed after $36\ h$. This comparison may be used to set a scale for signal *relevance*, used to reduce the number of genes to further analyze. Our analysis pipeline sets the scale for relevant alterations to be a 9/5-fold change in the Transcriptogram values, neglecting peaks with small differential Transcriptogram values in comparison to the most altered ones observed after $36\ h$. Naturally, when increasing the threshold required to select gene sets to be further analyzed, and hence decrease the number of false-positive errors, we will simultaneously increase the incidence of false-negative errors. We stress that the present analysis tries to address problems caused by the abundance of false-positive errors in finding biologically-relevant genes in experiments showing genome-wide transcription alterations.

Finally, we tested the gene sets in Clusters A, B, and C using *GSEA* using the *GSEA* default options (Signal-to-Noise, gene-set permutations). *GSEA* found these sets to be significantly differentially expressed after $30\,h$ after first RNA harvest (see also Table S 3 below), in agreement with our Transcriptogram results. *GSEA* defines a normalized enrichment score (NES) by normalizing ES (here the ratio between Transcriptogram values for *WT* and for *Mock* samples) by the expected value for a gene set with a given number of genes. Figure S 5 shows the evolution for each cluster of the average NES for each cluster follows the same trends as in Figure 5 in the main text.

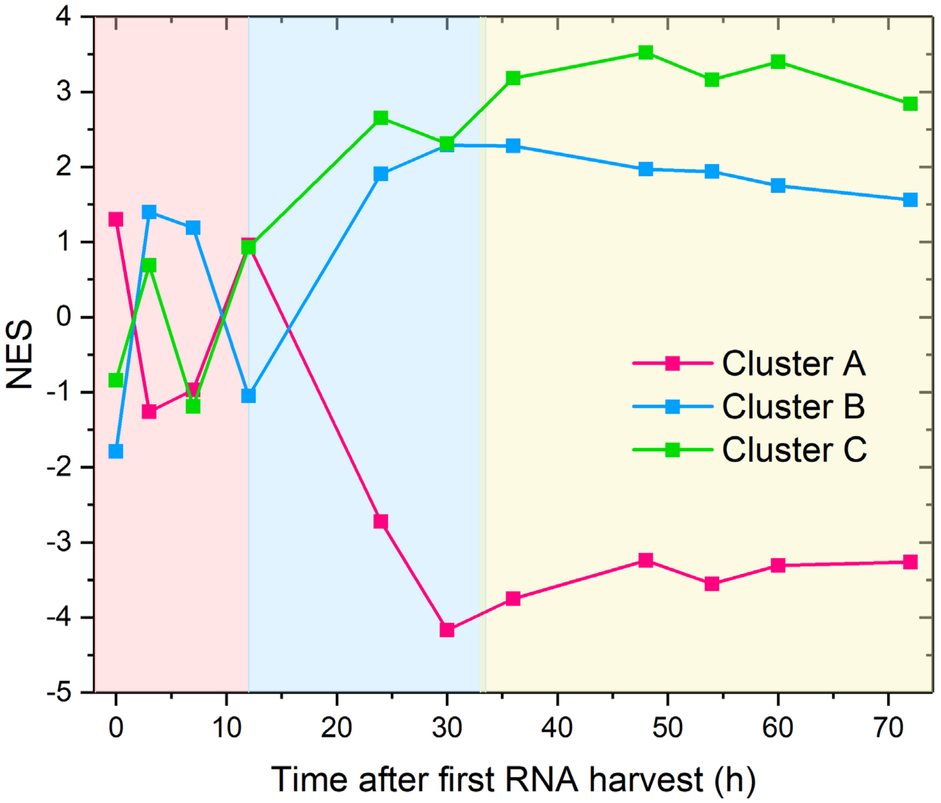


*Figure S 5. GSEA **NES** (Normalized Enrichment Score) using the GSEA options Signal-to-Noise statistics and gene-set permutation comparing WT and Mock samples. FDR <0.05 only after* $12\,h$ *after first RNA harvest (see Table S 2). After* $12\,h$ *the NES for each cluster follows the trends found using Transcriptograms.*

| Time after first RNA harvest $(h)$ | FDR | | |
|---|---|---|---|
| | A | B | C |
| 0 | 0.08 | 0 | 1 |
| 3 | 0.278 | 0.411 | 1 |
| 7 | 0.693 | 0.791 | 0.512 |
| 12 | 1 | 0.738 | 0.789 |
| 24 | 0 | 0.001 | 0 |
| 30 | 0 | 0 | 0 |
| 36 | 0 | 0 | 0 |
| 48 | 0 | 0 | 0 |
| 54 | 0 | 0 | 0 |
| 60 | 0 | 0.022 | 0 |
| 72 | 0 | 0.024 | 0 |

*Table S 3. False discovery rates (FDR) for the GSEA results for the gene sets represented by the A, B and C clusters found using the Transcriptogram method. Differential expression between Mock and WT samples is significant only after $12$ $h$ after the first RNA harvest.*

### 5. Complete panels for relative Transcriptograms and differential Transcriptograms

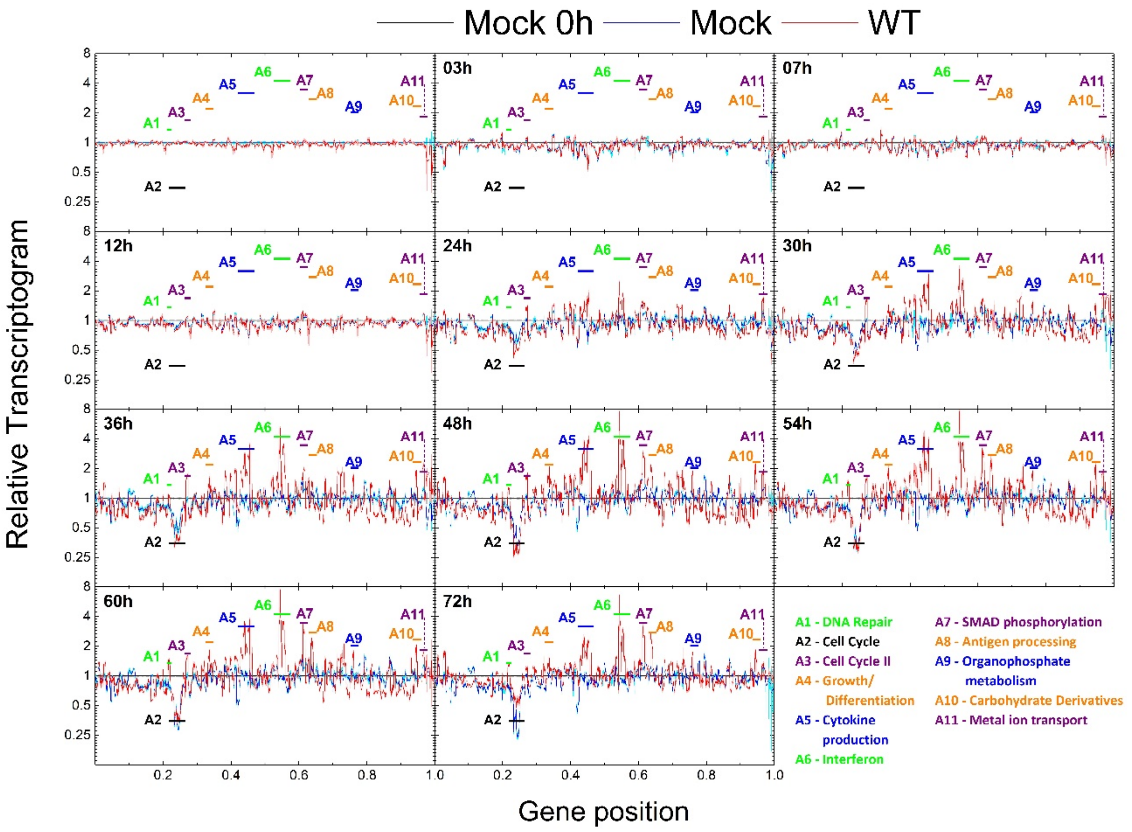


*Figure S 6. Relative Transcriptograms of radius 30 for Mock and WT samples at different timepoints after first RNA harvest, using the Mock sample at* $0\ h$ *as the control. Vertical axes are on* $log_2$ *scales. Black horizontal lines represent control (Mock) sample expression. Gray and light violet shading show the standard errors for the black and violet lines respectively. Major peaks are labeled with critical associated functions. The relative Transcriptograms for the Mock samples also change in time due to the effect of cell culture conditions on the cells.*

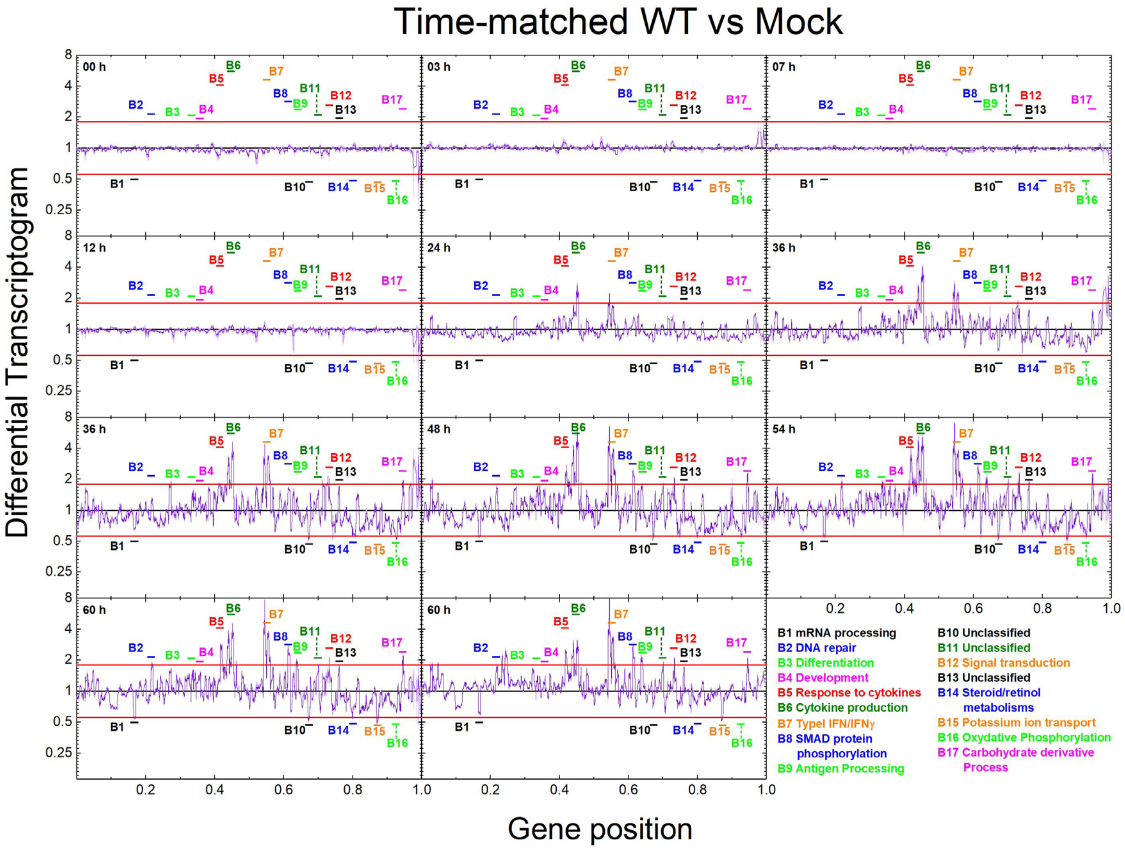


*Figure S 7. Differential Transcriptograms of radius 30 for WT samples, using the Mock expression at the corresponding time as the control. Times are measured from the first RNA harvest. Vertical axes are on $log_2$ scales. Black horizontal lines represent control (Mock) sample expression. Violet lines denote the differential Transcriptograms for WT samples. Gray and light violet shading denote standard errors for the black and violet lines, respectively. We identify 17 intervals where the violet line differs from control (Mock) by more than 1.8-fold at $54\ h$.*

### 6. Transcriptogram analyses for *DORF6* samples

In the original paper, the authors produced microarrays for Calu3 2B4 cells infected with a mutant strain of SARS-CoV with the ORF6 gene deleted (*DORF6*). ORF6 binds to karyopherin $\alpha 2$, trapping these proteins on intracellular membranes and sequestering karyopherin $\beta 1$. This sequestration may prevent nuclear import in general and hence act on transcription factors effect. The authors found that deletion of ORF6 attenuated the effects of viral infection both *in vitro* and *in vivo*, although viral titers were not significantly different from those for *WT* virus. The authors concluded that ORF6 tampers with innate immune signaling and other host signaling networks, to favor viral replication and pathogenesis.

This paper does not focus on the effect of deleting ORF6 in the virus may have on host cells, but on the transcription data for *WT* SARS-CoV immune response, as given by Figures 3 and 5 in the main text. Nevertheless, for completeness, we produced time-matched relative Transcriptograms for the *DORF6* transcription data series, using as controls the time-matched Transcriptograms for *Mock* samples, as shown in Figure S 8 . The peaks and valleys are essentially the same in the Transcriptograms comparing *WT* to *Mock* samples, with some subtle differences in intensity.

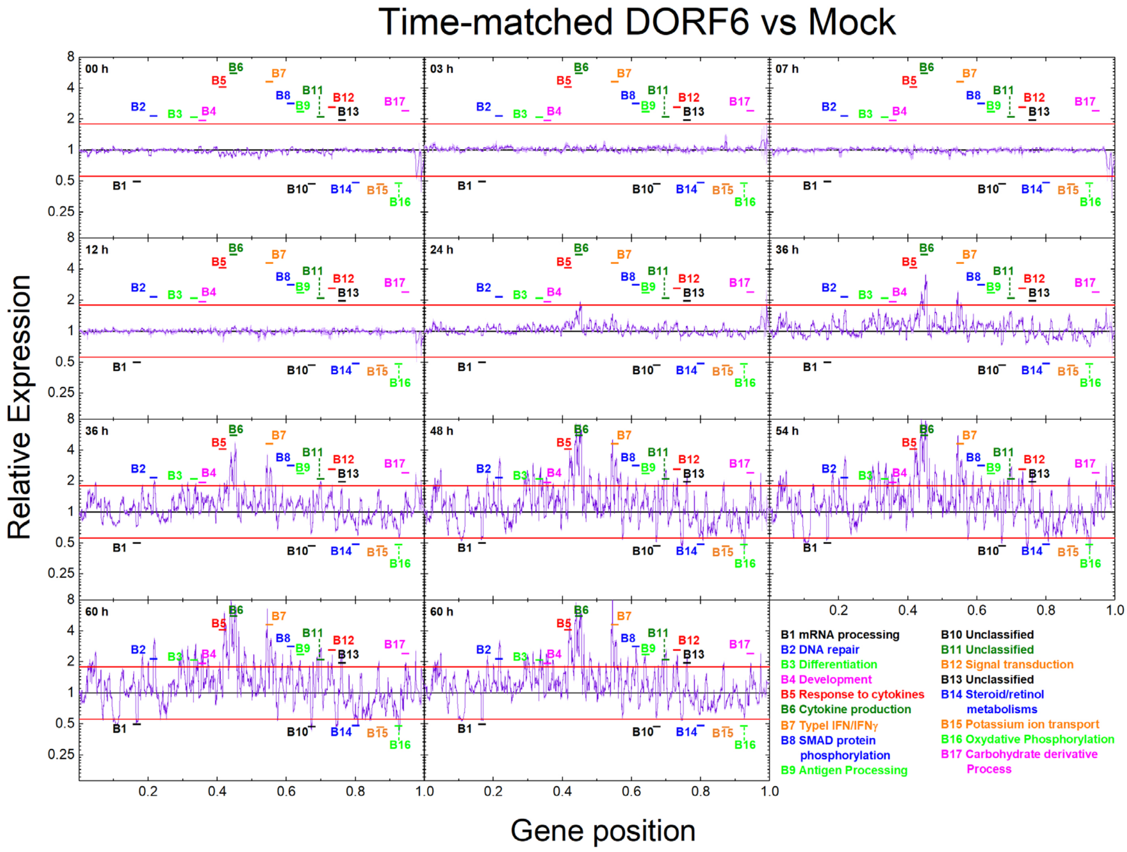


*Figure S 8. Differential Transcriptograms of radius 30 for DORF6 samples, using Mock expression at the corresponding time as control. Times are measured from the first RNA harvest. Vertical axes are on $log_2$ scales. Black horizontal lines represent control (Mock) sample expression. Violet lines denote the differential Transcriptograms for DORF6 samples. Gray and light violet shading denote standard errors for the black and violet lines, respectively. We identified the 17 intervals where the expression for WT samples were considered significantly different: DORF6 samples have essentially the same peaks.*

To highlight the differences between the expression changes in *WT* and *DORF6* samples, we produced time-matched relative Transcriptograms for *DORF6* using *WT* samples as controls, as shown in Figure S 9. Overall transcription differs between *WT* and *DORF6* after $48\ h$, but differences between *WT* and *DORF6* are much weaker than between *WT* and *Mock* or *DORF6* and *Mock*. In host cells ORF6 acts to prevent transcription factors from entering the nucleus and hence promote changes in gene expression. Transcription factors regulate many different pathways, affecting gene expression in a genome-wide manner. The differences we observe between *WT* and *DORF6*, though smaller, are statistically significant and genome wide, supporting the idea that the action of transcription factors produces the observed whole-cell phenotypic changes.

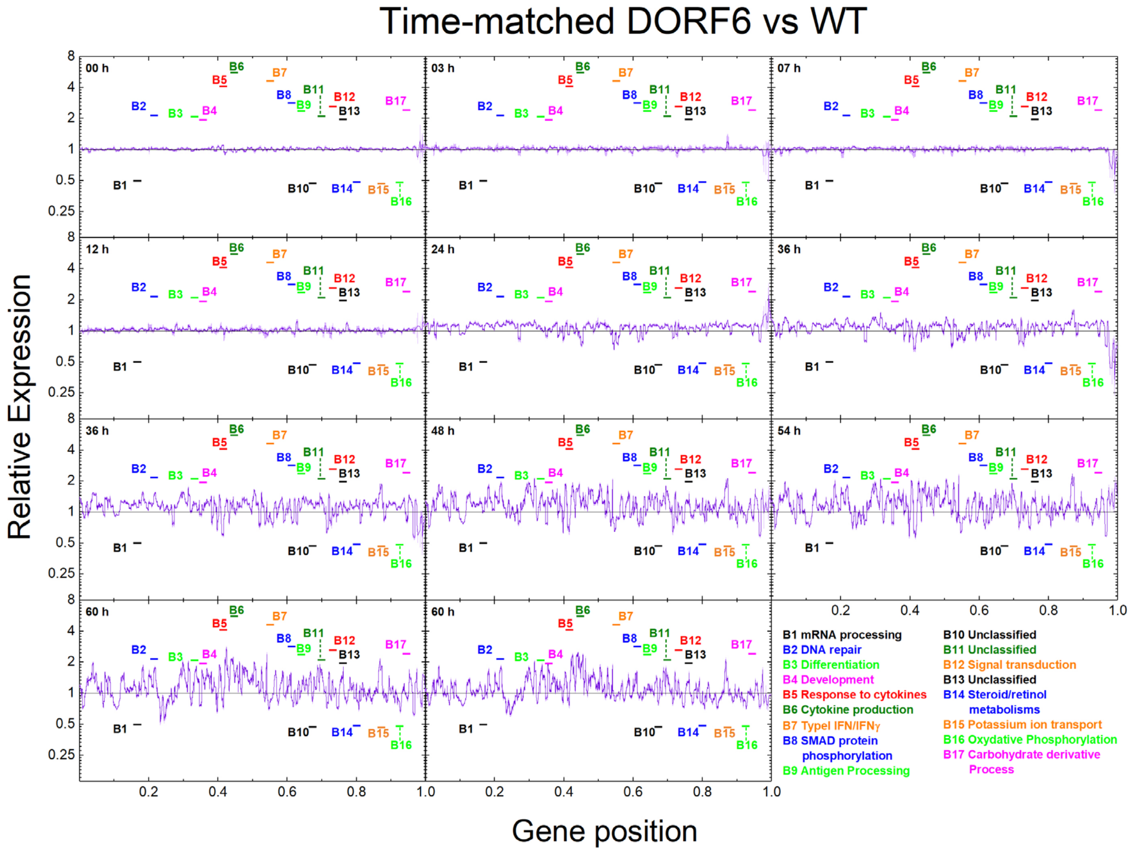


*Figure S 9. Differential Transcriptograms of radius 30 for DORF6 samples, using WT Transcriptograms at the corresponding times as controls. Times are measured from the first RNA harvest. Vertical axes are on $log_2$ scales. Black horizontal lines represent control (WT) sample expression. Violet lines denote the differential Transcriptograms for DORF6 samples. Gray and light violet shading denote standard errors for the black and violet lines, respectively. We have identified the 17 intervals where the expression of WT and DORF6 samples were considered significantly different from Mock samples.*

Complete analysis of Figure S 9 is beyond the scope of this paper. Sims and collaborators examined these differences in detail using other methods [1]. We examined expression changes for genes in the A, B and C clusters we identified using our Transcriptogram analysis for the *WT* virus strain. Figure S 10 reproduces relative expression averages for clusters A, B, and C for *DORF6* (bright colors) and *WT* (pale colors) data series. See Fig. 5 in the main text. Before 24 $h$ the *WT* and *DORF6* curves are similar for A, B and C. Between 24 $h$ and 72 $h$, cluster B and C *DORF6* curves grow steadily. The *WT* cluster B curve decreases after 30 $h$ and the *WT* cluster C curve decreases after 54 $h$. The difference between *DORF6* and *WT* cluster B time series is particularly dramatic, with *DORF6* levels significantly and persistentlyrita higher after 5 $h$. Since clusters B and C mainly comprise genes involved in immune response, we conclude that ORF6 affects the cellular immune response, in agreement with the original paper. The Transcriptogram analysis suggests that the most relevant differences in gene expression occur in genes in cluster B.

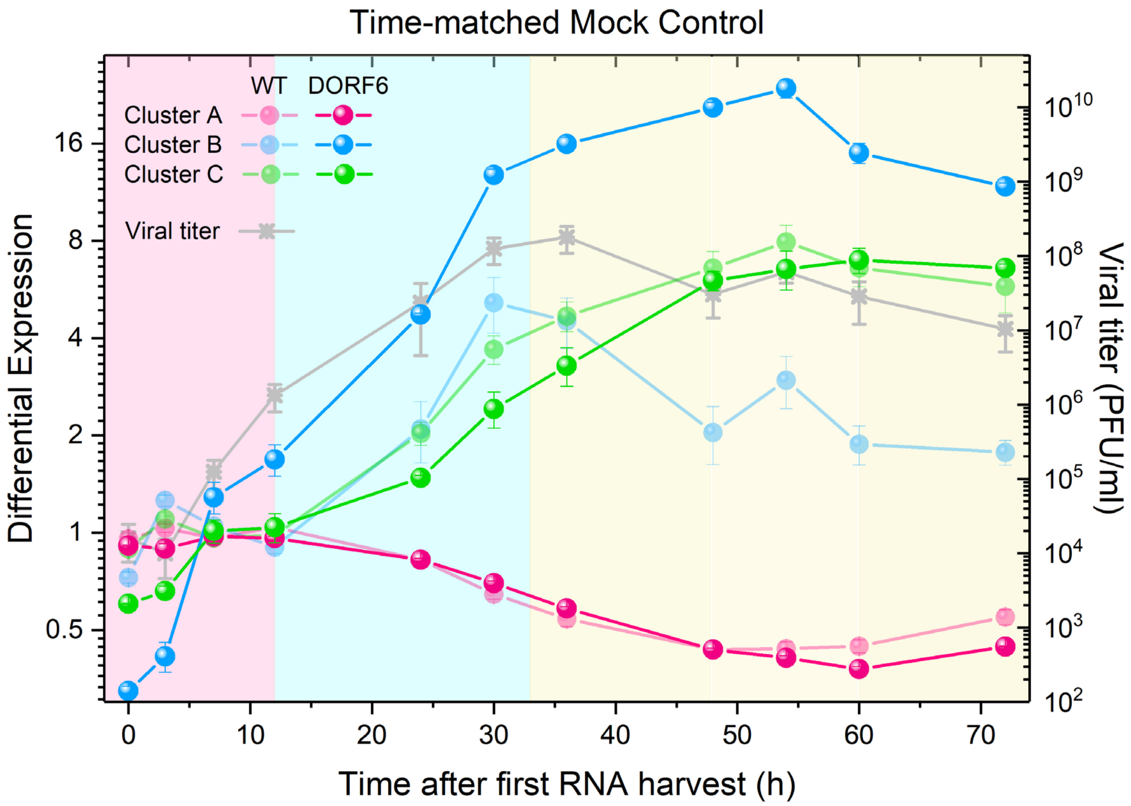


*Figure S 10. Time evolution of viral titer [1] (right $log_{10}$ axis) and average differential expression of the covariant gene clusters A, B and C (left $log_2$ axis) for DORF6 samples (intense colors) and WT samples (pale colors). The control for each time point is the time-matched Mock sample. Cluster A differential-expression dynamics are similar for DORF6 and WT. Cluster B responds very fast to viral infection, with DORF6 differential- expression below that of WT samples at $0\ h$, rapidly increasing to become much larger than for WT samples around $25\ h$. Cluster C differential-expression dynamics are similar for DORF6 and WT samples, with DORF6 levels slightly lower than WT between $20\ h$ and $55\ h$.*
